# Divergence and introgression among the *virilis* group of *Drosophila*

**DOI:** 10.1101/2022.01.11.475832

**Authors:** Leeban H. Yusuf, Venera Tyukmaeva, Anneli Hoikkala, Michael G. Ritchie

## Abstract

Speciation with gene flow is now widely regarded as common. However, the frequency of introgression between recently diverged species and the evolutionary consequences of gene flow are still poorly understood. The *virilis* group of *Drosophila* contains around a dozen species that are geographically widespread and show varying levels of pre-zygotic and post-zygotic isolation. Here, we utilize de novo genome assemblies and whole-genome sequencing data to resolve phylogenetic relationships and describe patterns of introgression and divergence across the group. We suggest that the *virilis* group consists of three, rather than the traditional two, subgroups. We found evidence of pervasive phylogenetic discordance caused by ancient introgression events between distant lineages within the group, and much more recent gene flow between closely-related species. When assessing patterns of genome-wide divergence in species pairs across the group, we found no consistent genomic evidence of a disproportionate role for the X chromosome. Some genes undergoing rapid sequence divergence across the group were involved in chemical communication and may be related to the evolution of sexual isolation. We suggest that gene flow between closely-related species has potentially had an impact on lineage-specific adaptation and the evolution of reproductive barriers. Our results show how ancient and recent introgression confuse phylogenetic reconstruction, and suggest that shared variation can facilitate adaptation and speciation.

## Introduction

Two major themes emerging from the rise of genomic approaches to phylogenetics and speciation are an understanding that genomic divergence is usually extremely patchy and that contemporary or historical introgression between species can be extensive. Heterogeneous genomic divergence has multiple potential causes. Genomic structure, especially inversions and other causes of variation in recombination rate, are associated with species’ divergence rates (Cruickshank and Hahn 2014; Wolf and Ellegren 2017). Natural and / or sexual selection may act locally on areas of genomes which therefore diverge more rapidly than the general ‘background’ genomic divergence rate (Nosil et al. 2009). Divergent regions probably contain barrier loci that contribute to adaptation and reproductive isolation, but genomic structure, demography and drift will also affect patterns of divergence, such that the interplay between these factors is complex (Ravinet et al. 2017). Gene flow following hybridisation may reduce divergence in regions of the genome, which successfully introgress between diverging species, and substantial proportions of such regions may be shared between species. There are now numerous examples of introgression between recent species, including during adaptive radiations (McGee et al. 2020).

The most notable example of gene flow between closely-related species is in hominins, where 1-3% of admixed proportion of DNA sequence in Eurasian populations results from introgression from Neanderthals (Sankararaman et al. 2014, 2016; Vernot and Akey 2014). *Heliconius* butterflies (Nadeau et al. 2013; Zhang et al. 2016; Edelman et al. 2019), African cichlids (Meier et al. 2017; Malinsky et al. 2018; Svardal et al. 2020), *Solanum* (Pease et al. 2016), Darwin’s finches (Lamichhaney et al. 2015, 2018; Han et al. 2017), *Anopheles* (Fontaine et al. 2015; Thawornwattana et al. 2018; Small et al. 2020) and *Drosophila* (Suvorov et al. 2021), all of which show evidence of substantial introgression. This may be more likely during recent divergence but in dire wolves (Perri et al. 2021), introgression seems to have occurred during early divergence within the clade but is absent between recent, isolated, species. Broad studies of introgression in *Drosophila* suggest it can be both ancient and recent (Suvorov et al. 2021). We do not know the extent to which interspecies hybridization may contribute to patchy genomic divergence; it would need to be extensive to cause background homogenization of the genome (as early verbal models suggested). Also, some chromosomes may be more resistant to introgression. Sex chromosomes often have disproportionate effects on reproductive isolation, so gene flow might be expected to be reduced on sex chromosomes compared to autosomes if they contain more barrier loci, resulting in higher levels of divergence on sex chromosomes (Ellegren et al. 2012; Wolf and Ellegren 2017). However, there are multiple reasons why sex chromosomes often show rapid divergence between species, including demographic effects and more effective background selection (Charlesworth et al. 1987).

The extent of introgression between species has revised our understanding of how gene flow may influence speciation. It has long been thought that considerable gene flow will restrict divergence between species, except in regions maintained by strong selection (Barton and Bengtsson 1986; Wu and Ting 2004). However, gene flow between lineages could also facilitate speciation by acting as a conduit for adaptive genetic variation (Marques et al. 2019). This ‘combinatorial’ view of speciation proposes that ancestral genetic variation can be reshuffled into unique combinations that may be favoured by ecological or other selection, circumventing the need for a build-up of de novo mutations. Long recognised in polyploid plants (Abbott et al. 2013), recently this has also been seen to occur much more extensively during homoploid speciation, including animals (Marques et al. 2019). Evidence of selection acting on introgressed genomic regions has been implicated in the maintenance of ecological barriers in *Heliconius* butterflies, Darwin’s finches, cichlids and sticklebacks (Nadeau et al. 2013; Zhang et al. 2016; Marques et al. 2016, 2019; Han et al. 2017; Meier et al. 2017; Samuk et al. 2017; Lamichhaney et al. 2018; Malinsky et al. 2018; Nelson and Cresko 2018; Svardal et al. 2020). However, quantifying the amount of introgression, adaptive or otherwise, during clade divergence remains an important challenge.

Species of the *melanogaster* group of *Drosophila* have arguably been studied most intensively. Genome scans have provided evidence for two patterns; inversions contribute disproportionately to genome-wide divergence, and introgression is lower on the X chromosome compared to the autosomes, as predicted if these contain more barrier loci (Garrigan et al. 2012; Lohse et al. 2015; Schrider et al. 2017; Turissini and Matute 2017; Mai et al. 2020; Korunes et al. 2021). In the *simulans* species complex, gene flow is extensive with 2.9% to 4.6% of the genome showing evidence of introgression between *D. sechellia* and *D. simulans*. These including genomic regions demonstrating selective sweeps suggesting adaptive introgression of genes involved in chemical perception (Garrigan et al. 2012; Brand et al. 2013; Schrider et al. 2017). Conversely, Turissini & Matute (2017) found minimal, older introgression between species in the *Drosophila yakuba* clade.

Here, we utilise new and existing sequence data from the *virilis* species group of *Drosophila* to examine patterns of genetic divergence and gene flow during divergence throughout the group. We have three main objectives. (1) to produce a timed estimate of the phylogeny of the group (2) to examine levels of divergence across autosomes and sex chromosomes and (3) to examine the extent and genomic location of introgression across the group, and to find out how large portion of it is potentially adaptive. Historically, the *virilis* group was thought to consist of 12 species that belong to two “phylads” or subgroups, the *montana* (usually thought to contain 8 species) or *virilis* phylads (5 species) (Throckmorton 1982). Species in the *virilis* group typically inhabit temperate ancient forest regions, with the exception of *Drosophila virilis* which is cosmopolitan and inhabits timber yards, breweries and market places (Patterson 1952; Throckmorton 1982). The group is thought to have originated in East Asia and spread into N. America via Beringia, and there are members of each phylad in the Nearctic and Palearctic regions (Throckmorton 1982). Morphological classification and molecular phylogenetics have failed to clarify the evolutionary relations of the group, especially the deeper branches (Patterson 1952; Chekunova et al. 2008; Morales-Hojas et al. 2011).

The *virilis* group has been used to study adaptation and speciation (Hoikkala and Poikela, in prep). Post-zygotic barriers, including sterility and inviability of the hybrids of both sexes, have evolved to varying degrees between species (Throckmorton 1982; Orr and Coyne 1989) and post-mating pre-zygotic (PMPZ) barriers have been shown to cause reductions in both interspecific and interpopulation fertilisation (Sweigart 2010; Ahmed-Braimah 2016; Garlovsky and Snook 2018; Poikela et al. 2019; Garlovsky et al. 2020). Species also show strong sexual isolation that is often asymmetric and based on species differences in male courtship song and / or cuticular hydrocarbons (Hoikkala and Lumme 1987; Liimatainen and Hoikkala 1998; Ritchie et al. 2001; Poikela et al. 2019). Also morphological differences, particularly in body pigmentation, have been documented in this group, though the evolutionary processes underpinning pigmentation differences is not yet well understood (Wittkopp et al. 2002, Wittkopp et al. 2002; Kulikov et al. 2004; Bubliy et al. 2007; Ahmed-Braimah and Sweigart 2015; Lamb et al. 2020). Several species of the *virilis* group, especially those of the *montana* phylad, persist in extreme cold environments, and show high cold acclimation and diapause which may contribute to genomic divergence (Vesala et al. 2012; Parker et al. 2015, 2016; Salminen et al. 2015; Tyukmaeva et al. 2015; Wiberg et al. 2021). Fixed and polymorphic chromosomal inversions have been reported within the group and may be driven by the activity and expansion of transposable elements, giving rise to regions of high differentiation between species and populations (Evgen’ev et al. 2000; Fonseca et al. 2013; Reis et al. 2018).

Here we use new whole-genomic sequence data from the *virilis* group to resolve their phylogeny and confirm that the major clade split is old and probably occurred in Miocene. We also show that gene flow has been extensive between some lineages despite the relatively rapid evolution of multiple sources of reproductive isolation between populations and species.

## Results

### Genome assembly

Altogether, we assembled 15 new genomes, with at least one for each of the species in the *Drosophila virilis* group, with the exception of *Drosophila texana*. Total genome size for the assembled genomes range from 170-210 MB, which corresponds to the range of published genomes of the *virilis* group. The new assemblies show relatively high completeness (>90%) using the BUSCO Diptera reference gene set (Supplementary table 1). Whilst most genes could be recovered, we found that higher levels of gene fragmentation, rather than missingness, accounted for lack of gene completeness, likely owing to limited genome contiguity. We identified 6-8% of the assemblies as repeat content, apart from *Drosophila ezoana* which showed higher levels (17%). With the exception of our annotation for *Drosophila lacicola*, we found between 13,100-18,075 genes in the genome assemblies, consistent with other Drosophila genome annotations. To supplement our analysis, we retrieved genomes for *Drosophila americana* (Fonseca et al. 2013), *Drosophila virilis* (Flybase v.1.07) and (as an outgroup) *Drosophila mojavensis* (Flybase v.1.04).

### Phylogenetic reconstruction and dating

We found agreement between the phylogenies produced by maximum likelihood and species tree reconstruction. All relationships within both the concatenated maximum-likelihood phylogeny and species tree were recovered with maximal (100%) bootstrap support. Our phylogeny is broadly consistent with previous phylogenetic reconstructions of the *virilis* group (Wang et al. 2006; Chekunova et al. 2008; Morales-Hojas et al. 2011), though we have better resolved the earlier branches, which influences interpretation of the group. In previous phylogenies (Orsini et al. 2004), *D. ezoana*, *D. kanekoi* and *D. littoralis* are included in the *montana* phylad, but our tree has the deepest branch separating these species, along with the *virilis* phylad, from the *montana* phylad. We suggest that the clearest resolution to this is to propose three phylads within the group: the *montana* phylad, containing *D. montana*, *D. lacicola*, *D. flavomontana* and *D. borealis*; the *virilis* phylad containing *D. virilis*, *D. lummei*, *D. novamexicana* and *D. americana* (and *D. texana*, not sampled here); and a *littoralis* phylad, containing *D. littoralis*, *D. ezoana* and *D. kanekoi* (Figure 1A). Within the *virilis* phylad, there is maximal support for divergence of *D. virilis* and *D. lummei* before the Nearctic *americana* clade, consistent with previous descriptions of the *virilis* phylad (Nurminsky et al. 1996; Caletka and McAllister 2004; Orsini et al. 2004; Wang et al. 2006; Chekunova et al. 2008; Morales-Hojas et al. 2011). Since no whole-genome data for *D. texana* is available, we could not fully resolve relationships within the *americana* clade. In comparison to the *virilis* phylad, there has been some disagreement regarding the placement of species within the *montana* phylad. In contrast to previous work, we show that species within the *montana* phylad are monophyletic with two sister lineages including the species pairs *D. borealis* and *D. flavomontana*, and *D. montana* and *D. lacicola*.

**Figure 1:**
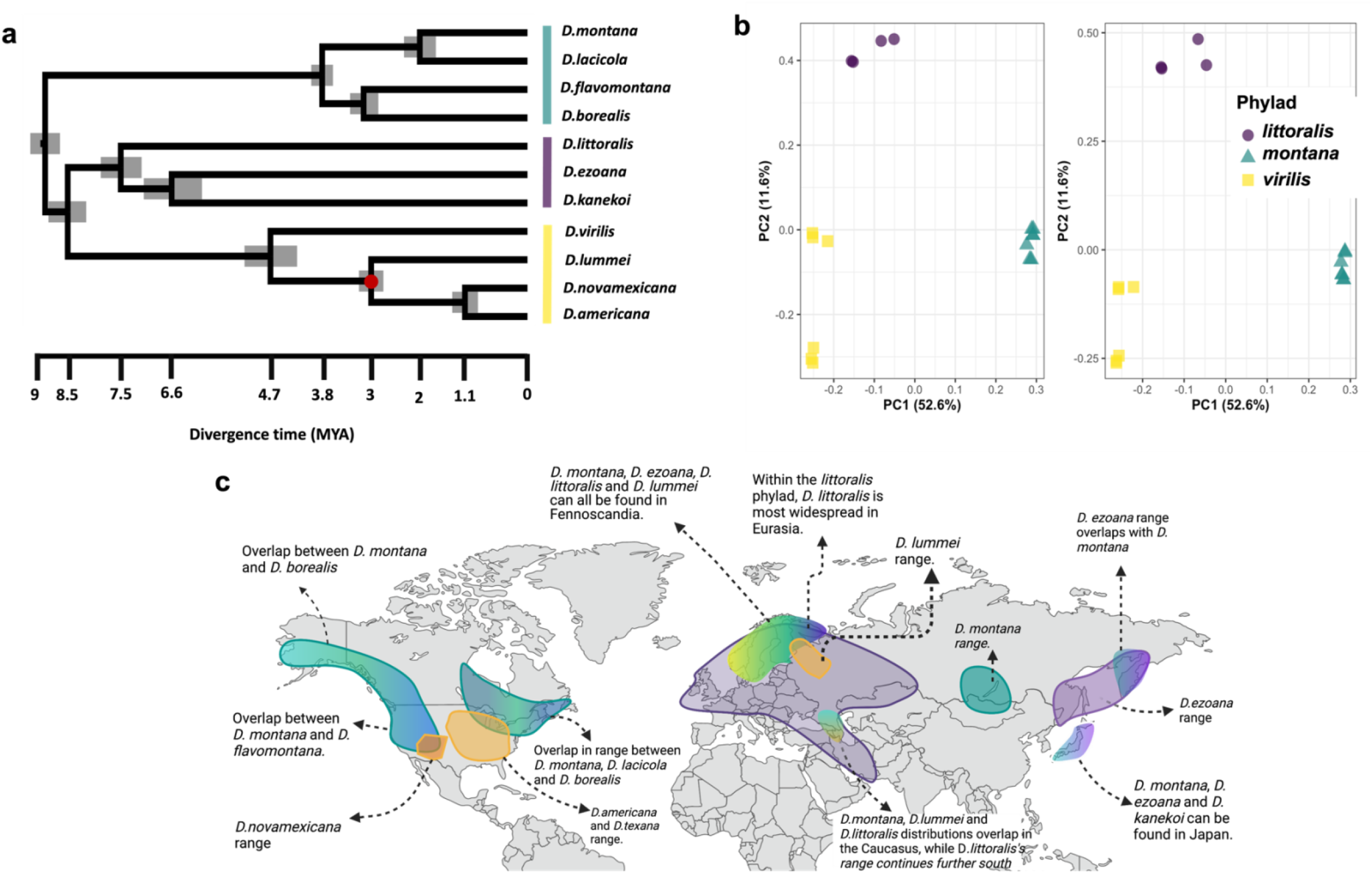
Species tree of the virilis group with estimated divergence times. **a)** Species tree reconstruction was performed using ASTRAL and gene trees for 1336 single-copy orthologs. Divergence times were estimated using BPP and randomly-sampled, genome-wide small introns (100 loci consisting of 75-85bp small introns). Posterior estimates for divergence times were scaled using a single calibration point (denoted by the red dot) set as 3.1-2.7 MYA at the split between *D. lummei* and *D. novamexicana* and *D. americana*, following rationale from Morales-Hojas et al. (2011). **b)** Principal component analysis (PCA) showing species relationships using randomly sampled SNPs on the X chromosome (3,218 SNPs) and autosomes (12,272 SNPs), respectively. **c)** Map showing putative ranges for species in all three phylads. Created with BioRender.com

We dated species divergence using putatively selectively neutral short introns (<80bp) randomly sampled genome-wide, with a single calibration point at the node characterising the split of the *americana* clade from *D. lummei* around 2.7-3.1 MYA, at the onset of the Northern Hemisphere Glaciation (Caletka and McAllister 2004; Morales-Hojas et al. 2011). Our inferred date for the basal node of the *virilis* group is 9 MYA, with the *littoralis* and *virilis* phylads diverging later on around 7.5 MYA (Figure 1A). Interestingly, the *littoralis* phylad radiated earliest and therefore contains the oldest species within the *virilis* group, with radiation of the *virilis* and *montana* phylads occurring 3.8 and 4.7 MYA respectively. Consistent with previous analyses, we show that divergence of the *americana* group within N. America occurred relatively recently (Caletka and McAllister 2004; Morales-Hojas et al. 2011). Phylogenetic relationships and the existence of three distinct clusters were also reflected across both the autosomes and the X chromosome in principal component analysis (PCA) (Figure 1B). Additionally, PCA showed that species within the *montana* phylad show tight clustering despite lineages diverging ∼4 MYA. Finally, we found no difference between mean divergence times estimated from coding regions in autosomes and the X chromosome (W = 42, p-value = 0.5)(Supplementary table 2).

### Introgression is pervasive between species of the *virilis* group

Whilst we recovered phylogenetic relationships with complete bootstrap support across the group, we note that relying solely on bootstraps to determine uncertainty in phylogenetic relationships may be misleading since maximal bootstrap support may often coincide with model misspecification (Yang and Zhu 2018), or considerable underlying gene tree conflict and systematic error (Kumar et al. 2012; Salichos and Rokas 2013). Using IQTREE2, we examined levels of gene and site discordance in gene trees used to infer the species tree. For every branch in the species tree, gene and site concordance factors are defined as the proportion of gene trees and sites in a given loci that are in agreement with the species tree. We found that branches leading to the *D. kanekoi* and *D. ezoana* species pair and the branch leading to *D. littoralis*, showed only 18% and 26% of decisive gene trees supported each respective branch in the species tree. Similarly, the same branches showed high levels of site-level phylogenetic discordance with around half the number of decisive sites (44% and 47%, respectively) found to be concordant with the species tree (Supplementary figure 1 & 2). Additionally, the branch leading to the *D. borealis* and *D. flavomontana* species pair also showed considerable levels of gene discordance (29%) and site level discordance (63%). In all three cases, the placement of these branches have been the most difficult to resolve in previous phylogenies (Morales-Hojas et al. 2011). The *virilis* phylad, on the other hand, had comparatively lower levels of gene tree discordance (Supplementary figure 1).

Whilst underlying gene tree conflict may be the consequence of poor phylogenetic signal or other technical issues associated with tree inference, it may also reflect genuine signals of gene flow or incomplete lineage sorting. To determine whether gene flow has occurred between species in the *virilis* group, we tested for excess shared, derived variants across the genome by calculating the minimum D-statistic (*Dmin*) for each possible trio in the group, where trios were conservatively organised in order to minimise the amount of introgression detected (Malinsky et al. 2020). We found that 35 of the 120 trios tested (35%) showed significant excess shared alleles after correcting for multiple testing, with mean excess allele sharing of 0.06 (6%). These results are incompatible with a single tree relating all species within the group and incomplete lineage sorting, and provides strong evidence for gene flow between some species within the group. Additionally, to supplement estimates of the minimum D-statistic, we calculated f4-ratios, which estimates the amount of ancestry in an admixed population that comes from potential donor populations. We find that only 12 trios show significant excess allele sharing (Dmin) and ancestry proportions (f4-ratio) over 1% (Figure 2). Of those 12 trios, introgression between *D. littoralis* and *D. ezoana* was consistently supported. We also found support for introgression between *D. lummei*, and *D. littoralis* and *D. ezoana*. Finally, we found consistent signals of introgression between *D. ezoana*, distributed in the northern parts of Europe, Asia, the Far East and in Japan, and the North American species of the *montana* and *virilis* phylad, which indicates that the ancestors of these species must have had overlapping distributions.

**Figure 2:**
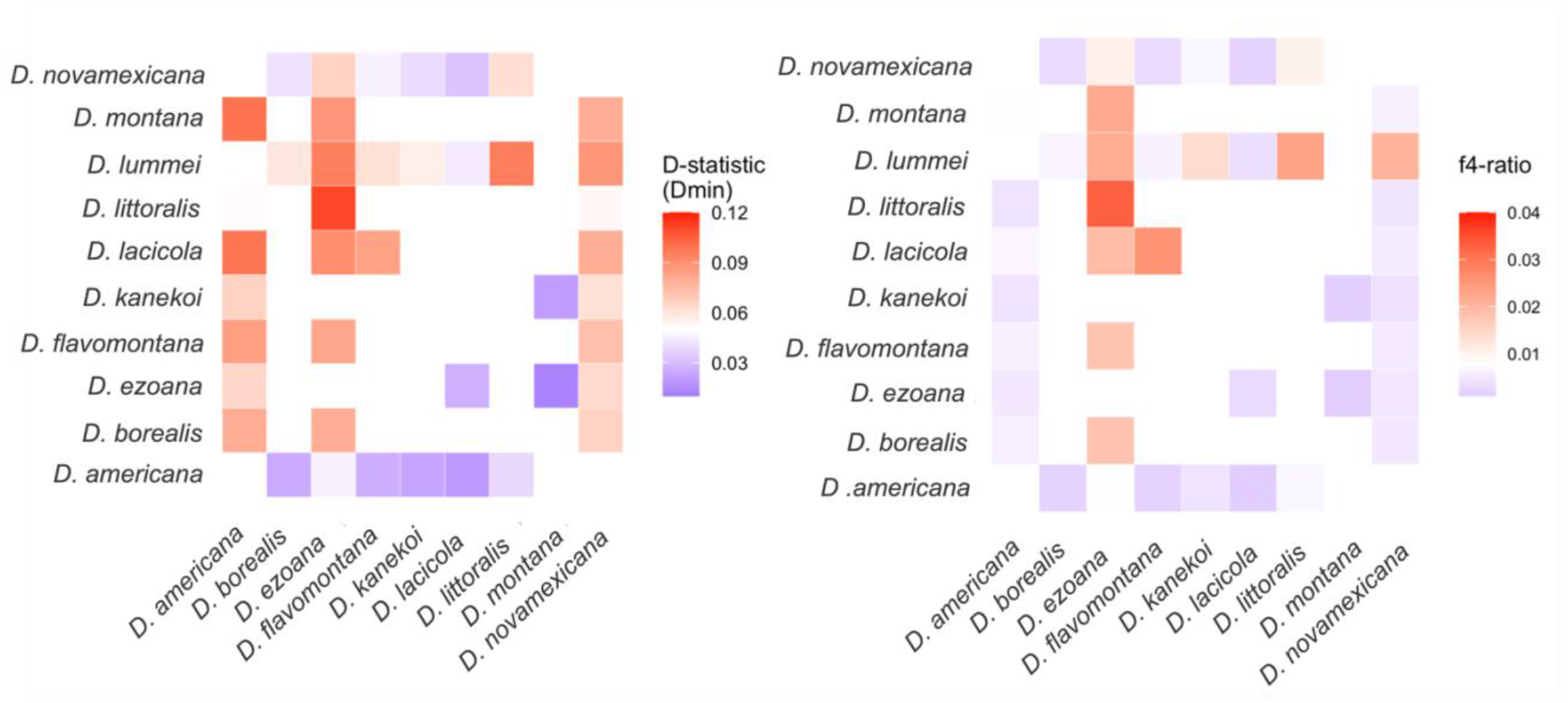
Gene flow is extensive between Palearctic and Nearctic members of the *montana* and *littoralis* phylads. On the left, D-statistic values shown between species with significant, excess allele sharing. On the right, admixture proportions (f4-ratio) between species with evidence of significant, excess allele sharing.

To investigate potential ancient introgression events, we calculated D-statistics by organising trios according to their species tree relationships, maximising the potential to detect gene flow events, as opposed to minimising the D-statistic (D_min_) to calculate conservative estimates of introgression. Using species tree relationships to organise trios, only 13 trios (out of 120 tested) showed significant D_min_ values exceeding 0.1 after correcting for multiple testing. These were mostly between species within the *montana* and *littoralis* phylads. To distinguish between individual gene flow events between multiple species, or a single ancestral gene-flow event affecting multiple descendant lineages, we calculated the f-branch metric (ƒ_b_ (C))(Malinsky et al. 2018). We found evidence for ancestral gene flow, with the branch leading to the *littoralis* phylad showing the highest level of introgression with *D. montana* (fb(C) = 28%; p < 0.001; Supplementary figure 3), and similarly high levels of introgression between the ancestral branch of the *littoralis* phylad and the other species of the *montana* phylad. This indicates that an ancestral gene flow event is at least partially responsible for the allele sharing between these groups of species.

Beyond genome-wide estimates of introgression, we tested for local phylogenetic discordance within phylads using TWISST. Such discordance can arise either due to gene flow or ILS. We found that the species tree was the most well-represented topology within each phylad. However, within the *montana* phylad, discordant topologies often showed comparable weighting to that concordant with the species tree across chromosomes (Figure 3A). This may suggest a lack of phylogenetic signal, consistent with incomplete lineage sorting genome-wide as well as heterogeneous patterns of gene flow. On the X chromosome, we found that a discordant topology with *D. borealis* and *D. lacicola* as this species pair had a higher average topology weighting than the species tree topology (ANOVA: df=2, F=1408.4, p < 0.001; Tukey multiple comparisons test: p < 0.001) suggesting extensive introgression on this chromosome. The other trios showed more consistent patterns of gene concordance across all chromosomes. Hence, it seems that interspecific gene flow (or ILS) may be more prominent in the *montana* phylad.

**Figure 3:**
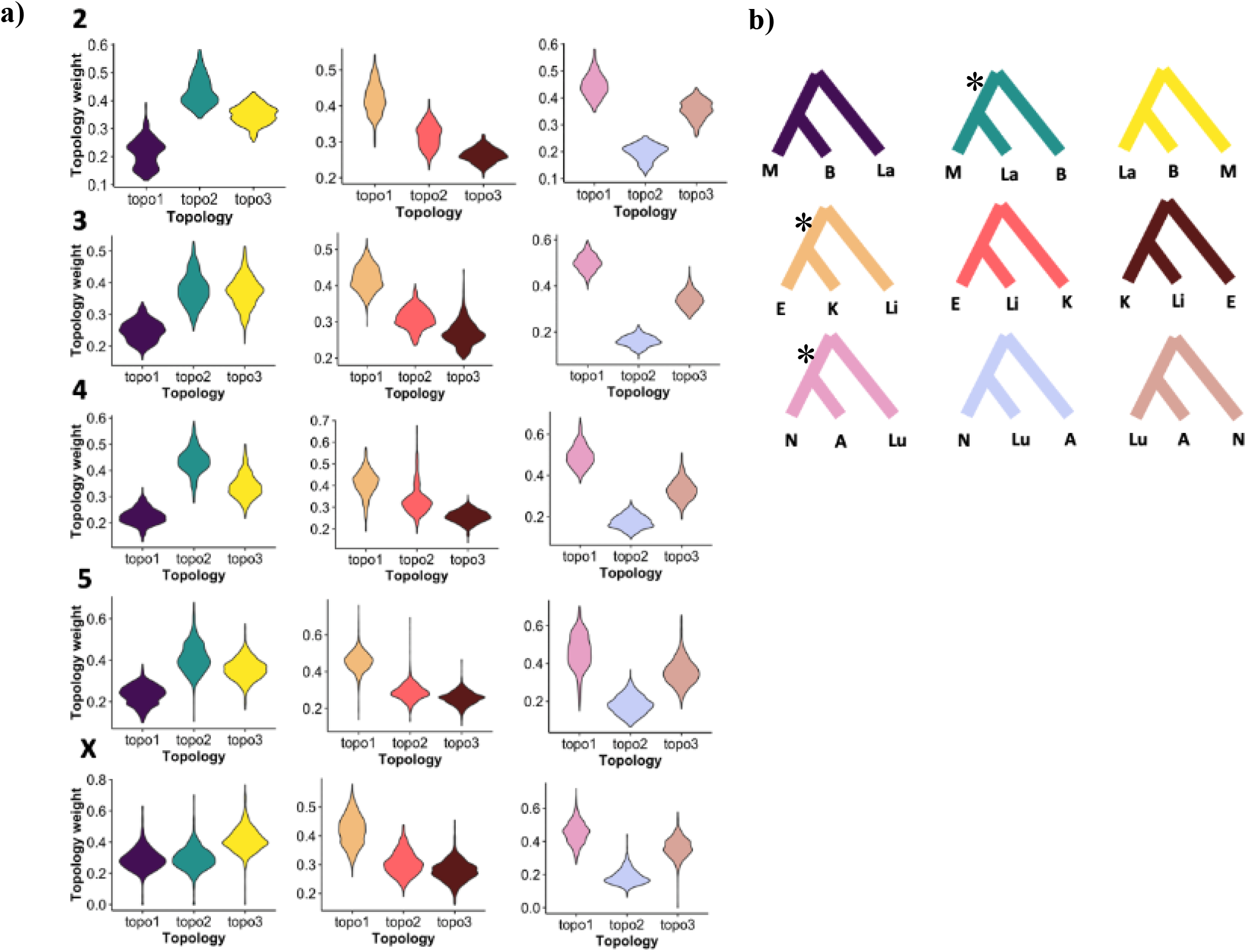
Topology weighting shows widespread phylogenetic discordance across the virilis group. **A)** Topology weighting for each possible topology split by phylad, where green, purple and yellow violins show alternative topology weightings for the *montana* phylad, orange, red and brown violins show alternative topology weightings for the *littoralis* phylad, and pink, light blue and tan trees show alternative topology weightings for the *virilis* phylad. Topologies with values of 1 indicate topologies with maximal weighting. Topology weighting separately for each chromosome, where each row is a different chromosome (numbers to the left of violin plots denote chromosome). **B)** Shows the topology each colour represents. Here, species names are abbreviated (M:*D. montana*, La:*D. lacicola*, B:*D. borealis*, F:*D. flavomontana*, E:*D. ezoana*, Li:*D. littoralis*, K:*D. kanekoi*, N:*D. novamexicana*, Lu:*D. lummei* and A:*D. americana.* Asterisks on topologies represent topologies that are concordant with the species tree.

### Localised patterns of genome-wide introgression

Additionally, we calculated genome-wide admixture proportions (f_dm_ and f_d_) in sliding windows. We found higher mean admixture proportions between species within the *montana* phylad (La <- F mean f_d_ = 0.04441; F <- M mean f_d_ = 0.04659; E <-Li mean f_d_ = 0.04065) compared to the *virilis* phylad (A <-Lu mean f_d_ = 0.02007), indicating evidence of recent admixture between sympatric species in the *montana* phylad. We note that in the *montana* phylad, the directionality of f_dm_ suggests considerable gene flow has occurred between *D. montana* and both *D. borealis* and *D. flavomontana* (Figure 5). In the *littoralis* phylad, we find that admixture has mostly occurred between *D. ezoana* and *D. littoralis,* with peaks of admixture localised to chromosome 4. Contrary to expectations, we find evidence of significantly higher admixture on the X chromosome between *D. lacicola* and *D. flavomontana*, when compared to autosomes (ANOVA: df=4, F=48.943, p < 0.001; Tukey multiple comparisons test: p < 0.001). This is not the case for other comparisons across the *virilis* phylad, where admixture proportions are usually considerably lower on the X chromosome compared to the autosomes (F <-M ANOVA: df=4, F= 6.5482, p < 0.001; E <- Li ANOVA: df=4, F= 64.584, p < 0.001)(Supplementary table 3).

To look for instances of overlap in admixture between the four trios tested, we extracted the admixture outlier windows (f_d,_ 95% quartile) for all trios and assessed overlap between these. We found little overlap between windows showing admixture proportions, with only one window showing overlap between two trios, and no windows showing overlap between more than two trios. To test what functions introgressed genes in each trio may contribute to, we performed gene ontology analysis for biological processes on genes found within the top 20 windows with the highest (10) and lowest (10) admixture proportions (f_dm_) for all trios. We found little overlap in the biological processes of genes exchanged between trios, but introgressed genes between *D. montana*, *D. borealis* and *D. flavomontana* showed significant (p = 4.98e-14) enrichment for heat shock proteins involved in polytene chromosome puffing and in insect stress responses (Zhao and Jones 2012) (Supplementary figure 4 & Supplementary table 4).

### Patterns of genome-wide divergence, gene flow and reproductive isolation

We calculated absolute divergence in coding and non-coding regions between species pairs to identify any differences between chromosomes, for example if ‘faster X’ divergence was seen. We found significant differences in absolute divergence between all chromosomes in non-coding and coding regions. However, inconsistent with expectations, we found only limited evidence for any faster X effect in species pairs across the *virilis* group in both coding and non-coding regions. Specifically, in only *D. montana* and *D. lacicola* (TMRCA: ∼2 Mya), and between *D. kanekoi* and *D. ezoana* (TMRCA: ∼6 Mya) we found significant, elevated divergence on the X chromosome relative to the autosomes (M-L ANOVA: df=4, F=218.41, p < 0.001; K-E ANOVA: df=4, F=59.086, p < 0.001) (Figure 4). Between *D. borealis* and *D. flavomontana,* and *D. novamexicana* and *D. americana*, we found the X chromosome (B-F mean X genic d_XY:_ 0.034; N-A mean X chromosome genic d_XY:_ 0.023) to show lower divergence compared to chromosome 4 (B-F mean chromosome 4 genic d_XY:_ 0.036) and chromosome 5 (N-A mean chromosome 5 genic d_XY:_ 0.024), respectively.

**Figure 4:**
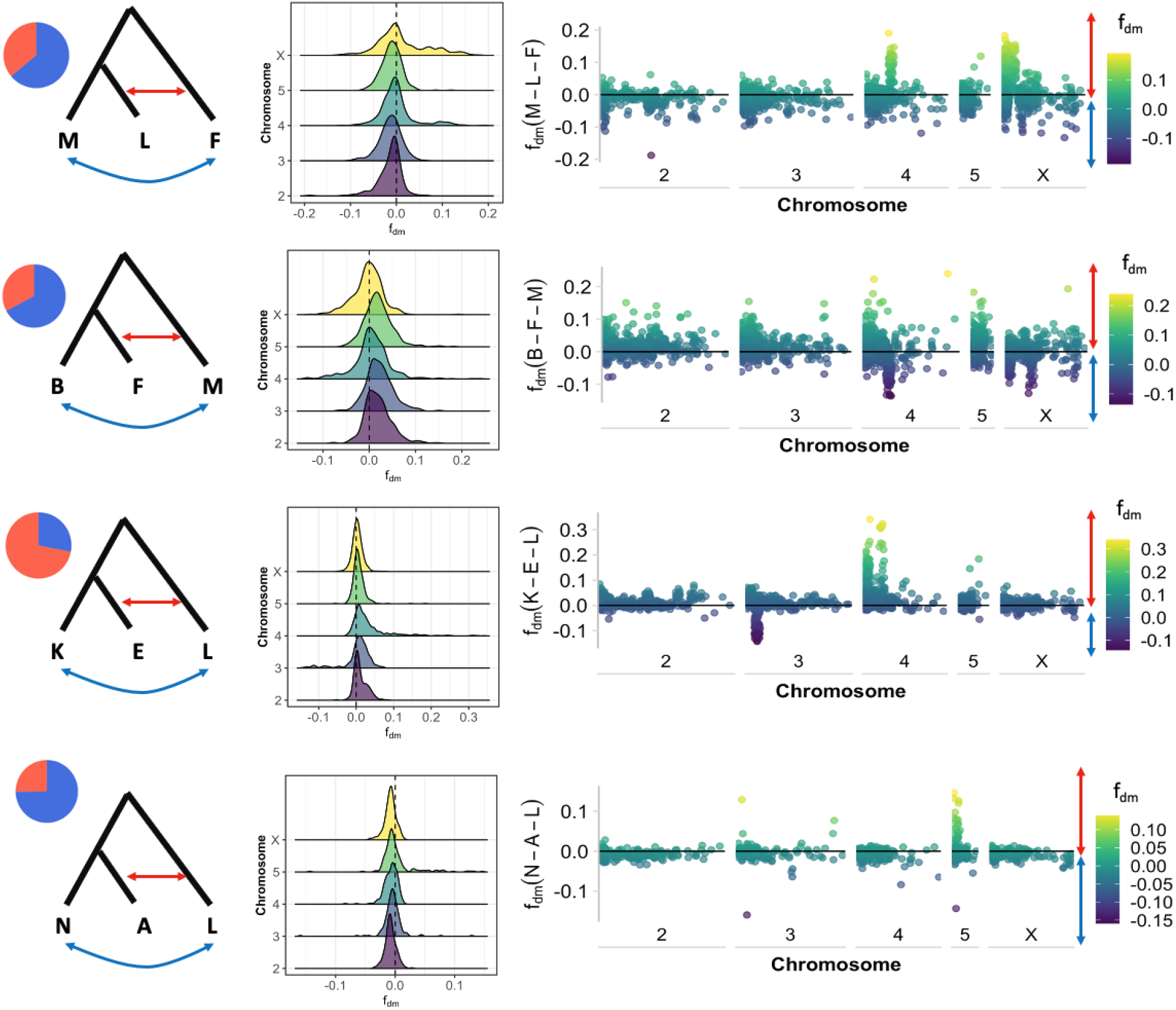
Genome-wide introgression facilitates phylad-specific gene sharing. **a)** Schematic showing directionality of gene flow tested by f_dm_ statistic and pie-chart illustrating proportion of introgression between species in each comparison calculated using f_dm_. **b)** Distributions of introgression across chromosomes for each comparison. Dotted line indicates neutrality (no allele sharing). **c)** Genome-wide introgression shown across chromosomes for each comparison, with solid line indicating neutrality.

**Figure 5:**
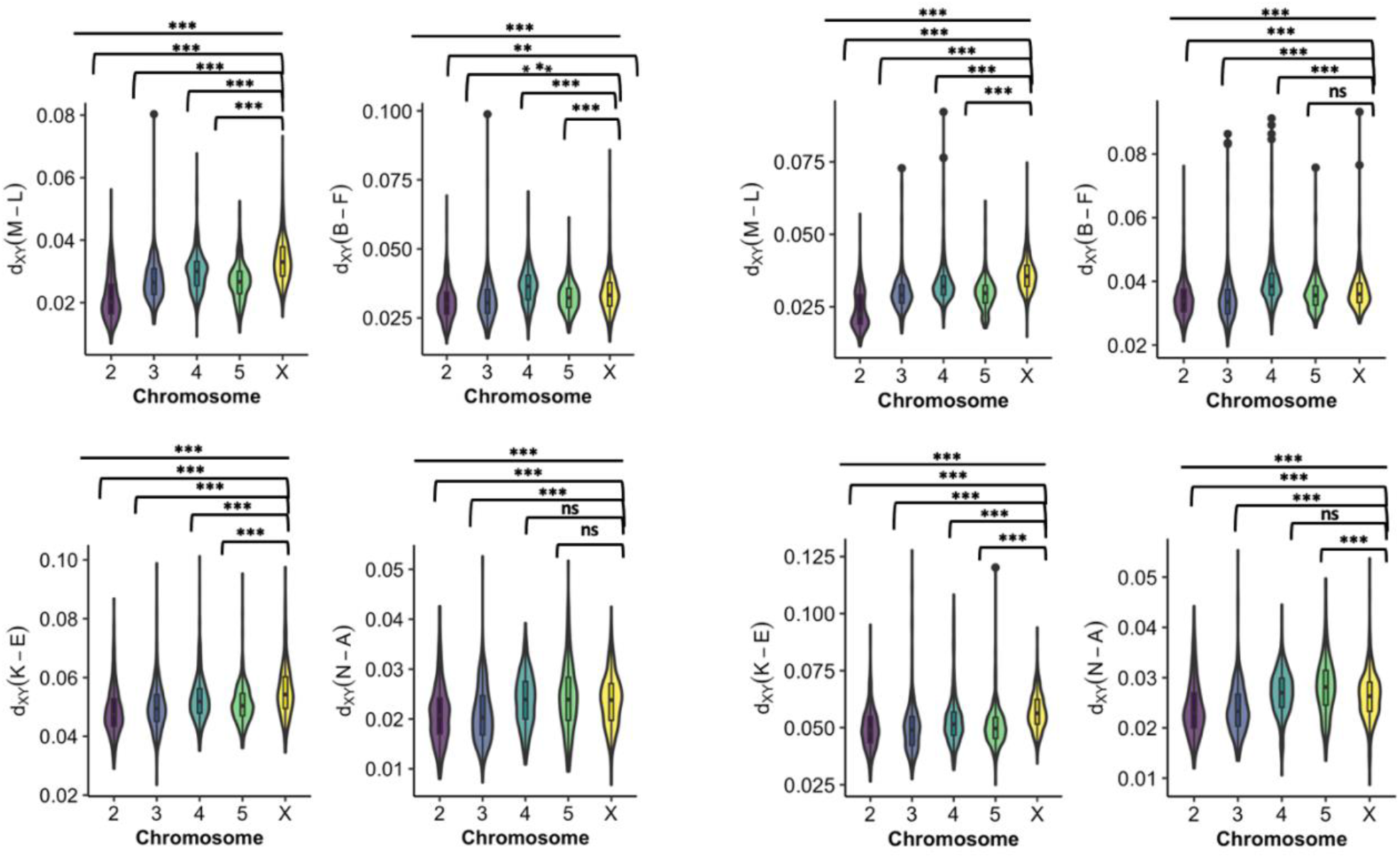
Absolute divergence calculated for each chromosome cross coding and non-coding regions. d_XY_ was calculated using genome-wide ∼50 kb windows between species pairs in the *virilis* group. **a)** On the left, d_XY_ was calculated for coding regions across the four species-pairs, and **b)** on the right, for intergenic regions. The species pairs shown here are the following: *D. montana* and D. *lacicola (M – L), D. borealis* and *D. flavomontana (B – F), D. kanekoi* and *D. ezoana (K - E) and D. novamexicana and D. americana (N – A*). Significance between all chromosomes was tested using an ANOVA and significance between the X chromosome and each of the autosomes was tested using the Tukey test. Stars indicate level of significance: 0 = ‘***’, 0.001 = ‘**’, 0.01 = ‘*’, 0.05 = ‘.’ and >0.05 = ‘ns’.

To understand broad patterns of reproductive isolation within the group, we collected data on pre-mating isolation and biogeography from Throckmorton (1982) and Yukilevich (2014), and paired this with genome-wide estimates of divergence (d_XY_) across the group. Specifically, previous comparative surveys of *Drosophila* have shown that pre-zygotic barriers generally evolve faster in sympatric species pairs compared to allopatric species pairs (Coyne and Orr 1989, 1997). After transforming estimates of d_XY_, pre-mating isolation and biogeography into dissimilarity matrices, we asked whether there was an association with biogeography – whether pairs share overlapping ranges (i.e. sympatry), or not (i.e. allopatry) – and estimates of pre-mating isolation. We found a significant positive correlation between biogeography and pre-mating isolation, whilst controlling for the effect of genome-wide divergence (Partial mantel test; R= 0.1823, p=0.016, permutations=10,000; Supplementary figure 5 and Supplementary table 5). In other words, sympatric pairs showed higher pre-mating isolation than allopatric pairs. Notably, three species comparisons (consisting of three closely-related species: *D. montana*, *D. borealis* and *D. lacicola*) show high levels of pre-mating isolation and relatively low genome-wide d_XY_ compared to most other species comparisons, indicating rapid evolution of pre-mating barriers.

### Molecular evolution of protein-coding genes across the *virilis* group

After correcting for multiple testing and filtering for saturation (dS > 2), we found 39 genes out of 7,443 genes with ω > 1 when calculating substitution rates across the entire *virilis* group (M0 model) (Figure 6 and Supplementary table 6). These included FASN2 (ω=1.08), a gene responsible for the production of methyl-branched cuticular hydrocarbons which contribute to reproductive isolation between *D. birchii* and *D. serrata* of the *montium* group (Chung et al. 2014) and are under sexual selection in *D. montana* (Veltsos et al. 2012; Jennings et al. 2014). Genes involved in sensory perception (Dhc36a, Or2a and CheA7A) were also among those showing evidence for rapid adaptive sequence evolution across the *virilis* group.

**Figure 6:**
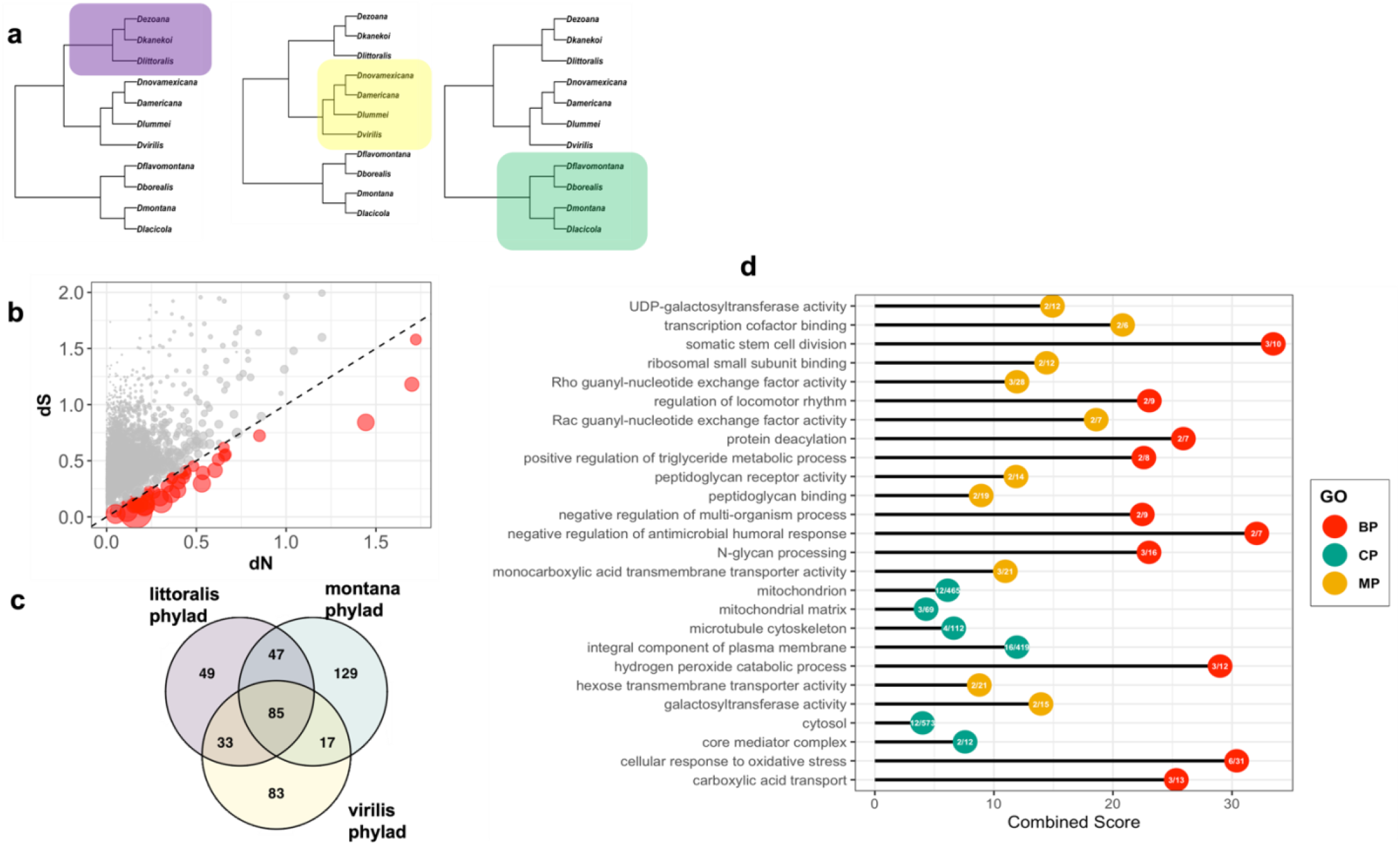
Detection of genes under putative positive selection in the virili*s* group. **a)** Illustration showing phylogenies and phylad tested in phylad-specific branch-site test using PAML. **b)** Synonymous substitution rate against non-synonymous substitution rate across entire gene tree for orthologs in the *virilis* group. Dashed line denotes dN/dS=1 and red circles are genes with dN/dS values exceeding 1. Size of circles corresponds to dN/dS ratio **c)** Overlap in genes found in branch-site test between the three phylads tested. **d)** Gene ontology for genes detected to be under selection during phylad-specific branch-site test. Gene ontology terms in legend are: Biological Process (BP); Cellular Process (CP) and Molecular Process (MP). Inside each circle is the number of genes under putative selection that overlap with the reference gene set for each gene ontology term. Only terms with a significant p-value included and the top ten ontology terms for each category shown (except for CP, where only 6 genes showed significant p-values).

Finally, we conducted phylad-specific branch-site tests, with branches in each phylad considered foreground branches with different rates of ω to background branches (bsC model vs. M1) (Figure 6a). We corrected for multiple testing (FDR=0.01) and filtered for saturation (dS > 2, dN/dS > 2), and detected 214 genes in the *littoralis* phylad, 278 genes in the *montana* phylad, and finally, 218 genes in the *virilis* phylad, that had significantly different branch-site ratios compared to background branches (Supplementary Table 7). When comparing overlap between these three comparisons, we found 83 genes with significantly different branch-site ratios across all three phylad-specific tests. Between the *littoralis* and *montana* phylads, and between the *littoralis* phylad and the *virilis* phylad, we found an overlap of 47 and 33 genes respectively, with the *montana* phylad and the *virilis* phylad only sharing an overlap of 17 genes. Gene ontology inference of biological processes related to genes detected using phylad-specific branch-site tests indicated genes found in phylad-specific tests are involved in regulation of locomotor rhythm, antimicrobial response and regulation of triglyceride metabolic process.

## Discussion

A major challenge in speciation genomics is the identification of patterns of genetic variation associated with the emergence and maintenance of reproductive barriers. One way of addressing this is by characterising levels of genetic divergence, gene flow and the strength of reproductive barriers in species pairs across the speciation continuum. For example, a comparative survey examining rates of migration and genomic divergence across animals found ‘a grey zone of speciation’, where at 0.5% - 2% net divergence a cessation of gene flow was observed (Roux et al. 2016). Collectively, these analyses have yielded important, general observations: (a) genome-wide effective migration rate reduces with higher levels of genetic divergence, (b) heterogenous patterns of genome-wide genetic divergence are expected to be the combined result of gene flow, divergent selection and genomic features such as recombination rate, (c) sex chromosomes show higher genetic divergence compared to autosomes, and (d) sexual isolation is generally higher in sympatric species pairs compared to allopatric species pairs (Coyne and Orr 1989, 1997). In Drosophila, broad-scale comparative analyses have so far been limited to single species pairs or complexes, with few exceptions (Mai et al. 2020; Suvorov et al. 2021). Using de novo whole-genome data and genome assemblies, we examined the prevalence of phylogenetic discordance, gene flow, levels of genome-wide divergence and measures of reproductive isolation in the Drosophila *virilis* group.

Phylogenetic placements in the *virilis* group have been the source of some contention. Previous inferred phylogenies have suggested that *D. littoralis, D. kanekoi* and *D. ezoana* are more closely-related to the *montana* phylad than to the *virilis* phylad (Orsini et al. 2004; Andrianov et al. 2010). In agreement with Morales-Hojas et al. (2011), we found that these species are more closely-related to the *virilis* phylad, but are quite distinct, and we suggest that they represent an additional phylad within the group, termed *littoralis* after the oldest species. We also resolved species pairs within the *montana* phylad. Importantly, we show that the difficulty in resolving the phylogeny has likely been the result of pervasive gene flow between closely-related species within the same phylads, and ancient introgression between the common ancestors of the phylads. Using a combination of D-statistics and gene and site-concordance factors, we detected evidence of strong phylogenetic incongruence in branches leading to the *littoralis* phylad. Between closely-related species, we also found considerable heterogeneity in genome-wide phylogenetic incongruence likely attributable to more recent gene flow between species, particularly in the *montana* phylad where species often share overlapping geographic ranges (Figure 1c).

We quantified divergence times for species in the group using putatively selectively-neutral small intronic regions (Haddrill et al. 2005; Halligan and Keightley 2006). In general, our estimates of species divergence fit with estimates of divergence times from previous analyses (Morales-Hojas et al. 2011). The oldest radiation within the group is that of the *littoralis* phylad, which is ∼7.5 MYO. Little is known about the ecology and distribution of *D. kanekoi*, except that its range is limited and largely endemic to Japan, alongside more widespread species in the group like *D. montana*, *D. virilis* and *D. ezoana*. Considerably more is known about the ecology of *D. littoralis* and *D. ezoana* that are found in northern Finland together with *D. montana* and *D. lummei* (Aspi et al. 1993). The divergence of the *montana* and *virilis* phylads occurred later, around the beginning of the Pliocene, where temperatures likely warmed and passage into North America via the Bering Strait was possible (Marincovich and Gladenkov 1999; Robinson et al. 2008). Divergence of both *D. montana* and *D. lacicola*, and *D. borealis* and *D. flavomontana* in the *montana* phylad roughly coincide with the split of *D. lummei* from the *americana* clade in the virilis phylad and the intensification of the Northern Hemisphere Glaciation (Caletka and McAllister 2004; Bartoli et al. 2005; Ruggieri et al. 2009). Altogether, we suggest that changes in climate during the Pliocene may have facilitated the spread and early divergence of both the *virilis* and *montana* phylads.

The phylogeographic origin of the *montana* phylad, and in particular *D. montana*, has been the subject of recent debate. Unlike other members of the *montana* phylad, whose ranges are restricted to N.America, *D. montana* co-exists in parapatry with *D. flavomontana and D. borealis* in N.America, overlaps with *D. ezoana* and *D. littoralis* in Eurasia and with *D. ezoana* and *D. kanekoi* in Japan (Figure 1). Our results indicate that the divergence of *D. montana* and *D. lacicola* occurred around ∼2 MYA, though little is known about the range and ecology of *D. lacicola*, so it is difficult to determine what may have driven this split. Recent modelling of divergence time between *D. montana* and *D. flavomontana* (Poikela et al, in prep) suggested considerably earlier divergence times (1-2 MYA) for the divergence of the *montana* phylad than previous phylogenies and recent demographic analysis (Morales-Hojas et al. 2011; this study). These slight differences in divergence times likely reflect differences in parameter scaling and genomic partitions chosen for analysis. As a result, interpretations of biogeographical scenarios associated with the split of the *montana* phylad are varied. For example, earlier estimates are consistent with divergence of *D. montana* and the *montana* phylad in N.America (Poikela et al, in prep), whilst older estimates may indicate a potential origin for *D. montana* in Asia (Mirol et al. 2007; Morales-Hojas et al. 2011; Garlovsky et al. 2020). However, given the range of *D. montana,* it is possible that the ancestral lineage of the *montana* phylad had a Holarctic distribution leading to vicariant speciation events in N.America.

Interspecific hybridization is common across closely-related species, but the question of when gene flow is expected to cease between distantly-related species is still unclear. We calculated genome-wide estimates of gene flow using D-statistics and found only 10% of tested trios to show any evidence of gene flow. However, after accounting for phylogeny and non-independence of gene flow estimates using the f-branch statistic (Malinsky et al. 2020), we only found strong evidence for an ancient gene flow event(s) between the ancestor of *montana* phylad and the *littoralis* phylad lineage, prior to the divergence of *montana* phylad. We found no evidence for independent gene flow events between *D. montana* and *D. littoralis, D. ezoana* and *D. kanekoi*, suggesting either (a) that strong barriers to gene flow arose quickly in Eurasian populations of *D. montana* or (b) that the lack of gene flow is indicative of divergence of the *montana* phylad in N.America. These analyses demonstrate the difficulty in disentangling recent hybridization events from ancient hybridization events across relatively speciose taxonomic groups (Malinsky et al. 2018; Ferreira et al. 2020).

Interpreting patterns of genetic divergence is difficult due to the effect of recombination rate and linked selection (Noor and Bennett 2009; Cruickshank and Hahn 2014). Whilst absolute divergence is not affected by within-population diversity and linked selection, it is expected to be affected by recombination rate variation and gene flow, such that divergence is lower in regions of low recombination and gene flow can homogenise peaks of divergence (Nachman and Payseur 2012). Here, we observed heterogenous patterns of divergence across ‘true’ species pairs, and in striking contrast with previous analyses across diverse taxonomic groups (Meisel and Connallon 2013; Charlesworth et al. 2018), we found unexpected patterns of low genetic divergence on the X chromosome compared to autosomes in two of the four species pairs. Interestingly, in the *montana* phylad we observed evidence for a faster X chromosome effect between *D. montana* and *D. lacicola*, but not between *D. borealis* and *D. flavomontana*. One potential explanation for these patterns are differences in rates, direction and timing of gene flow between species in the *montana* phylad. Whilst little is known about the range of *D. lacicola*, our local introgression analysis (f_dm_) suggest most introgression has occurred recently between *D. montana*, *D. borealis* and *D. flavomontana,* with some evidence of comparable levels of shared variation on the X chromosome and autosomes. But there has been extensive introgression between the X chromosomes of *D. borealis* and *D. lacicola*. Additionally, our data does not allow us to characterise or describe inversions despite their prevalence in the *virilis* group and their potential contribution to patterns of genomic divergence and speciation (Reis et al. 2018, 2020).

As with previous studies, we also observed a positive correlation between admixture proportion and absolute divergence across species pairs and genomic regions (Supplementary figure 6)(Stankowski et al. 2019; Duranton et al. 2020). In European lineages of sea bass, archaic genomic tracts were shown to contribute to excess absolute divergence and windows with high admixture proportions were also predicted to be involved in reproductive isolation (Duranton et al. 2020). Whilst introgressed variation may be contributing to genome-wide divergence in the *virilis* group, it is difficult to determine what forces may be responsible for the correlated genomic landscape observed here without information on recombination rate variation.

The role of introgression on adaptation and speciation has received considerable attention. In some cases, the transfer of beneficial alleles into recipient taxa can facilitate adaptation (Whitney et al. 2006, 2010; Dasmahapatra et al. 2012; Racimo et al. 2016; Malinsky et al. 2018; Oziolor et al. 2019; Valencia-Montoya et al. 2020). The exchange of locally adaptive introgressed variation can also contribute to speciation whilst globally adaptive introgressed variation cannot (Abbott et al. 2013). However, reliably identifying adaptive introgression remains challenging. We found heterogenous patterns of introgression genome-wide, with peaks of introgression specific to each compared trio. Additionally, functions of genes found within high-confidence introgressed windows were trio-specific, even between closely-related species in the *montana* phylad, suggesting independent gene flow events and/or selective purging of introgressed variation following introgression events.

Reproductive barriers between species in the *virilis* group have been well-characterised and supplemented broad comparative surveys of reproductive isolation and genetic divergence in *Drosophila* (Throckmorton 1982; Coyne and Orr 1989, 1997; Yukilevich 2014). Here, we recapitulate some of the patterns observed before by demonstrating an association between biogeography and pre-mating isolation, where sympatric species comparisons generally show higher levels of pre-mating isolation compared to allopatric species comparisons. In particular, most species in the *montana* phylad show strong pre-mating isolation despite relatively low genome-wide divergence and evidence of pervasive gene flow, indicating a complex demographic history and possible reinforcement of existing pre-mating barriers. This is supported by recent work showing almost complete reproductive isolation between *D. montana* females and *D. flavomontana* males with some evidence supporting reinforcement of sexual and postmating-prezygotic barriers in the reciprocal cross in sympatric populations, as well as cascade reinforcement of these barriers between *D. flavomontana* populations (Poikela et al. 2019). Altogether, patterns of genetic variation within the *montana* phylad likely reflect older gene flow events that may have contributed to reproductive isolation.

## Conclusion

The *virilis* group diverged relatively rapidly and evolved varied, but strong isolation mechanisms in the face of gene flow. Ancient gene flow between the *montana* and *littoralis* phylads likely confused previous attempts to fully reconstruct the speciation history of the group. Within the *montana* phylad, introgression has been extensive, likely reflecting a recent history of gene flow between closely-related species. We suggest that genes evolving rapidly throughout the group may play a role in sexual isolation and that introgressed variation is possibly adaptive. We draw from previous analyses of reproductive isolation within the group and find evidence that gene flow may have contributed to reinforcement within the *montana* phylad. Differences in genetic divergence across chromosomes do not clearly support a disproportionate role for sex chromosome in facilitating isolation between species pairs in the *virilis* group. Our study clarifies the phylogenetic relationships between species in the *virilis* group and highlights the potential for gene flow to shape evolutionary history and patterns of genetic variation genome-wide.

## Methods

### Stocks; obtaining and sequencing samples

For genome sequencing we used a single individual flies (female or male) which were derived from isofemale strains collected at different locations or received from the stock centers (for more details see Supplementary table 1). DNA was extracted with CTAB method followed by RNAse treatment and multiple phenol-chlorophorm-isoamyl alcohol and chlorophorm-isoamyl alcohol cleaning steps. The integrity of the extracted DNA was assessed with agarose gel and the quantity was measured with qubit (Thermo Fischer Scientific). The samples were shipped on ice to Edinburgh Genomics, UK, for library preparation and genome sequencing.

### Genome assembly and annotation

The samples were prepared using Nextera libraries protocol and sequenced on Illumina HiSeq 2500 platform (150bp paired-end reads). The read quality was checked with fastqc (Andrews 2010) and the assemblies were done with MaSuRCa genome assembler (Zimin et al. 2013). Genome quality was assessed using BUSCO (v.2.0) with the Diptera gene set. To annotate the genomes we used BRAKER (v.2.1.4) with GenomeThreader to map D. *virilis* homologs, which was used as evidence, and subsequently used to train ab initio gene prediction via AUGUSTUS. We then filtered predicted genes using AGAT program ‘agat_sp_filter_incomplete_gene_coding_models.pl’ (https://github.com/NBISweden/AGAT) for genes without a start and stop codon.

### Read mapping, variant calling and filtering

FASTQ reads were first trimmed using Trimmomatic (v0.36) with the following parameters (ILLUMINACLIP:TruSeq3-SE:2:30:10LEADING:3TRAILING:3SLIDINGWINDOW:4:15 MINLEN:36). Reads for all samples, including the D. *virilis* and D. americana genomes obtained from Flybase, were mapped to a PacBio reference genome (Poikkela et al, in prep) using bwa-mem (0.7.15). We used samtools (v.1.6) to sort, mark and remove duplicate reads, and subsequently, we used to GATK Genome Analysis tool to realign reads around indels. We called variant and invariant sites using bcftools (v.1.6) and filtered sites using bcftools for depth (DP>10) and mapping quality (MQ>30).

### Resolving the phylogenetic tree and divergence times

To investigate phylogenetic relationships within the *virilis* group, we utilised proteins characterised as complete and single-copy for each species using BUSCO. For species with multiple de-novo genomes, we selected protein sets with the highest BUSCO completeness percentage. After retrieving BUSCO protein sets for each species, we used Orthofinder (v2.3.12) to identify orthologs from species-specific protein sets. Multiple sequence alignments of single-copy orthologs were retrieved from Orthofinder and subsequently aligned using MAFFT (v7.147b). We then filtered gappy and poorly-aligned columns from all alignments using trimAl (v1.4.rev15) with -gappyout option. In total, 1336 alignments remained and were used for phylogenetic analysis. We inferred a maximum likelihood phylogeny using a concatenated alignment of these single-copy orthologs and a JTT+F+R5 model determined by ModelFinder (Kalyaanamoorthy et al. 2017) and 1000 bootstraps in IQTREE (v.2.0.3). Concatenation of alignments produced 787, 425 bp sequences for all 12 species, with 18,827 parsimony-informative sites and 64,533 singleton sites. Since concatenating loci can result in failure to resolve the species tree (Mendes and Hahn 2018), we also inferred individual gene trees using maximum likelihood in IQTREE2 for each gene alignment and subsequently used these for species tree reconstruction in ASTRAL (v5.6.3). Given a set of gene trees, ASTRAL attempts to construct a species tree from the maximum number of quartet trees represented in all given gene trees, thereby accounting for possible gene tree discordance. In both the concatenated maximum-likelihood phylogeny and species tree, *D. mojavensis* was used as the outgroup. Finally, to assess phylogenetic concordance amongst the gene trees, we estimated gene concordance and site concordance factors using IQTREE. Orthofinder was performed on BUSCO single-copy protein sequences for each genome to retrieve orthogroups.

To date divergence times and infer ancestral population size for each node, we first randomly extracted small introns (<80bp) from the VCF file using bedtools (v2.29.2) and VCFtools (0.1.14). Here, small introns were chosen due to previous work in *Drosophila* showing selective constraint in small introns was comparable to selective constraint in selectively neutral, four-fold degenerate sites. Extracted small introns were then concatenated into small blocks of 300bp based on their proximity to one another. In total, we used 100 regions around 300bp long. We then converted these 300bp regions into multiple sequence alignments with the requirement that all species contain a SNP for each column in the alignment, using the script vcf2phylip.py. We estimated divergence time and ancestral population size under the multiple-species coalescent model using BPP (v.4.1.4) (Flouri et al. 2018). Specifically, we use the A00 model which estimates divergence time and population sizes using a fixed species tree and provided species-to-sample relationships. For this, we provided the species tree inferred by ASTRAL. We then converted divergence time estimates from coalescent units into real dates using a single calibration point at the node separating the Eurasian *D. lummei* from the American (*D. americana* and *D. novamexicana*) species. Following previous analysis, we assume the last common ancestor of *D. lummei* and the ‘*americana*’ clade had a Holarctic distribution and diverged following the Northern Hemisphere Glaciation event estimated to have occurred around 2.7-3.1 Mya (Caletka and McAllister 2004; Morales-Hojas et al. 2011). For calibration and estimation of lower and upper-bound divergence, we used the bppr package in R.

Additionally, to identify differences in divergence and ancestral population size between autosomes and the X chromosome, we randomly extracted coding regions in the autosomes and X chromosomes separately, and produced 100 loci each consisting of 300bp coding sequences for autosomes and X chromosomes. Again, we used vcf2phylip.py to produce multiple sequence alignments containing columns where all samples contained SNPs – columns with missing data for any sample were discarded for each loci. We then estimated divergence times and ancestral population sizes for autosomes and the X chromosome using the A00 model in BPP.

Finally, we used principal component analysis to assess whether genome-wide variation supported species relationships found using orthologs. We extracted coding regions from the autosome and X chromosome separately and pruned SNPs using bcftools ‘+prune’ command with a window size of 100. We then used PLINK2 to further prune the dataset down using windows of 50kb, step sizes of 10 and r^2^ threshold of 0.1 (Purcell et al. 2007). For the X chromosome, 3,218 variants were used for the PCA, and for the autosomes 12,272 variants were used.

### Estimating gene flow

To test for introgression we calculated excess allele sharing using D-statistics (Green et al. 2010) using filtered variants (described above). Specifically, we calculated D_min_, defined as the minimum amount of allele sharing regardless of any assumptions made about the tree topology and species relationships, for each trio in the *virilis* group. Additionally, to more accurately determine D_min_, we provided the species tree constructed using ASTRAL as an input species tree. We corrected for multiple testing using a Bonferroni correction and *D* values with P < 0.5 were considered significant. Additionally, to determine when introgression may have taken place, we calculated the f-branch statistic (Malinsky et al. 2018, 2020). The f-branch statistic (ƒ_b_ (C)) measures admixture between a donor species C and branch *b* by calculating admixture for all possible combinations of ƒ(A,B,C;O), where A and B are sister lineages, and A is a descendant of branch *a* and B is a descendant of branch *b.* In short, The f-branch statistic (ƒ_b_ (C)) calculates the amount admixture that has occurred between taxon C and the branch (*b*) leading to a descendant taxon B, relative to the admixture that has occurred between taxon C and a sister branch to taxon B (branch *a*), where A is a descendant of branch *a* and B is a descendant of branch *b (*ƒ(A,B,C;O)). Significant f-branch (ƒ_b_) values indicate ancient introgression between branch *b* and taxon C. Admixture proportions were also calculated in windows using the f_d_ (Martin et al. 2015) and f_dm_ (Malinsky et al. 2015) statistic. Both statistics were calculated in windows of 100bp with a step size of 200bp.

To supplement this, we constructed genome-wide phylogenies and weighted topologies using TWISST (Martin and Belleghem 2017). Phylogenies were calculated in three clades: the *virilis* phylad, the *littoralis* phylad and the *montana* phylad. Phylogenies were constructed from filtered variants (described above) using the GTR model in PhyML in windows of 100bp. TWISST analyses were performed using the ‘complete’ parameter method, calculating exact topological weightings by considering all possible sub-trees. For all phylogenies, *D. virilis* was used as the outgroup.

Additionally, we converted the filtered VCF file into a genotype-specific file and then used popgenWindows.py to calculate genetic divergence (d_XY_) in 50kb windows for all possible species pairs (https://github.com/simonhmartin/genomics_general). To aid in Coyne & Orr-style (1989, 1997) comparative analysis, we retrieved data on pre-mating isolation and biogeography for species pairs in the group from (Yukilevich 2014), and where biogeography data was missing, we used biogeography inferences from Throckmorton (1982) to supplement the analysis.

### Molecular evolution across the *virilis* group

Gene models produced by annotation were extracted from a representative genome for each species in the *virilis* group. Orthofinder was used to cluster orthologs (Emms and Kelly 2015, 2019). Multiple sequence alignments were extracted from Orthofinder output, and putative orthologs were filtered for paralogs by removing species with more than one sequence in every orthogroup. A custom script was used to retrieve corresponding nucleotide sequences based on amino acid alignments of orthologs produced by Orthofinder. Alignments were then filtered for length, removing alignments with sequences smaller than 150 nucleotides. Additionally, alignments with less than 8 species were filtered at this stage too. Alignments were then made using MAFFT with default options (Katoh and Standley 2013). Finally, alignments were trimmed using trimAl with the parameter ‘--gappyout’ (Capella-Gutiérrez et al. 2009). Species trees for each file were constructed for each alignment by pruning species from the species tree that were filtered from the alignments. Here, we first assessed d*N*/d*S* across the *virilis* group and for each alignment using the M0 model via ete3 (Yang et al. 2000; Huerta-Cepas et al. 2016).

Additionally, we used a clade-specific branch-site test to identify genes evolving rapidly on each of the three clades in the *virilis* group using a clade-specific branch-site test (bsC vs. M1)(Yang and Nielsen 2002). Models were compared using a likelihood ratio test and p-values were corrected for multiple testing strictly using false discovery rate (p < 0.01). To understand the function of significant genes we used Blast2GO (Conesa et al. 2005). To obtain more specific gene ontology predictions, we retrieved one representative sequence from the alignments of significant genes and blasted them against NCBI non-redundant database. Uniprot identifiers for best-hits were then converted into corresponding D.melanogaster orthologs, where possible, using Flymine (Lyne et al. 2007). Finally, we used FlyEnrichr for gene annotation and gene ontology prediction (Kuleshov et al. 2019).

## Acknowledgments

This study was primarily supported by an NERC (UK) award, NE/J020818/1, and associated NBAF award NBAF852. LY was supported by a University of St Andrews studentship. We thank Svetlana Sorokina from the Koltzov Institute of Developmental Biology, RAS, for help with the fly collection, valuable comments to the manuscript and providing the *D. littoralis* AB10-51 line. Additionally, we would like thank the staff of ‘Raduga’ biological station for their help in our fly collection in Kamchatka, Russia. We are grateful to Stephen Goodwin, Megan Neville, Konrad Lohse and Darren Parker for advice and support at various stages of the project.

## Data accessibility

Whole-genome sequencing reads are available at the NCBI Sequence Read Archive under BioProject PRXXXXXXXXX. Scripts for analysis will be available at: https://github.com/LeebanY/Virilis_Comparative_genomics.

## Author contributions

MGR and AH conceived the study; VT performed field- and lab-work; LY analysed the data, with contributions from VT (genome assembly); LY wrote the manuscript with input from MGR, AH and VT.

---

**Supplementary figure 1:**
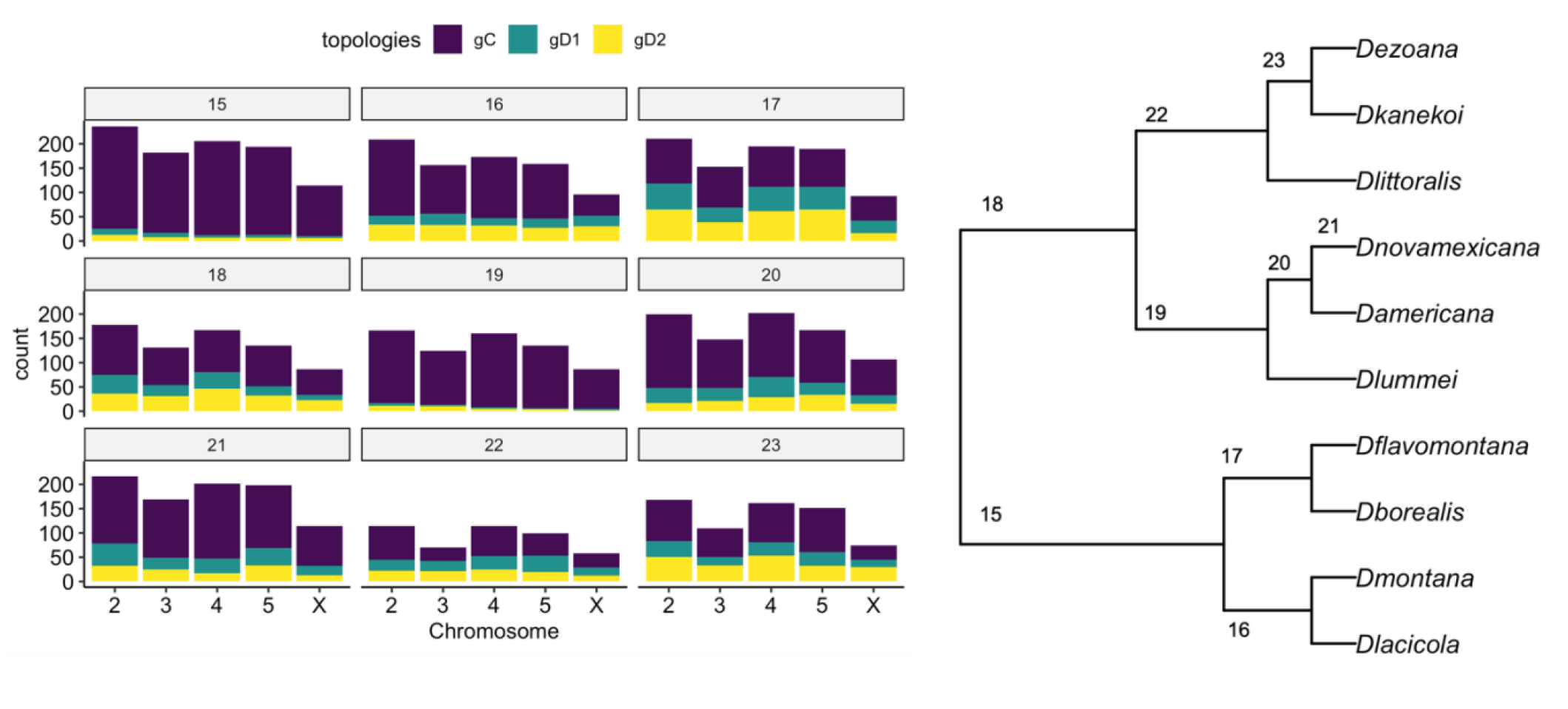
Gene concordance across nodes and chromosomes on the species tree. On the y-axis, counts denote the number of decisive gene trees supporting the species tree topology (gC; purple) and alternative topologies (gD1 and gD2; green and yellow). Number labels on the top of each plot show gene concordance for specific nodes on the species tree phylogeny (indicated on the left).

**Supplementary figure 2:**
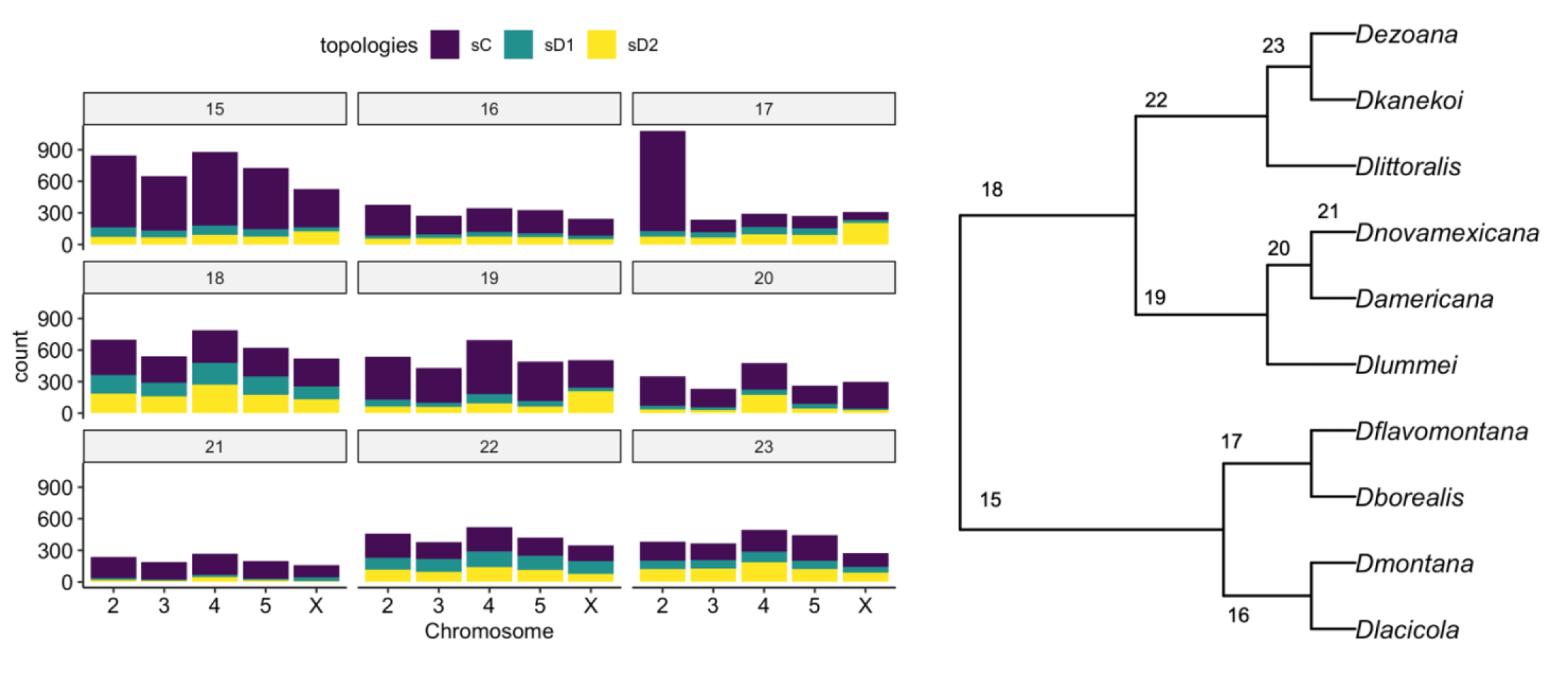
Site concordance across nodes and chromosomes on the species tree. On the y-axis, counts denote the number of decisive sites in the alignment supporting the species tree topology (sC; purple) and alternative topologies (sD1 and sD2; green and yellow). Number labels on the top of each plot show gene concordance for specific nodes on the species tree phylogeny (indicated on the left).

**Supplementary figure 3:**
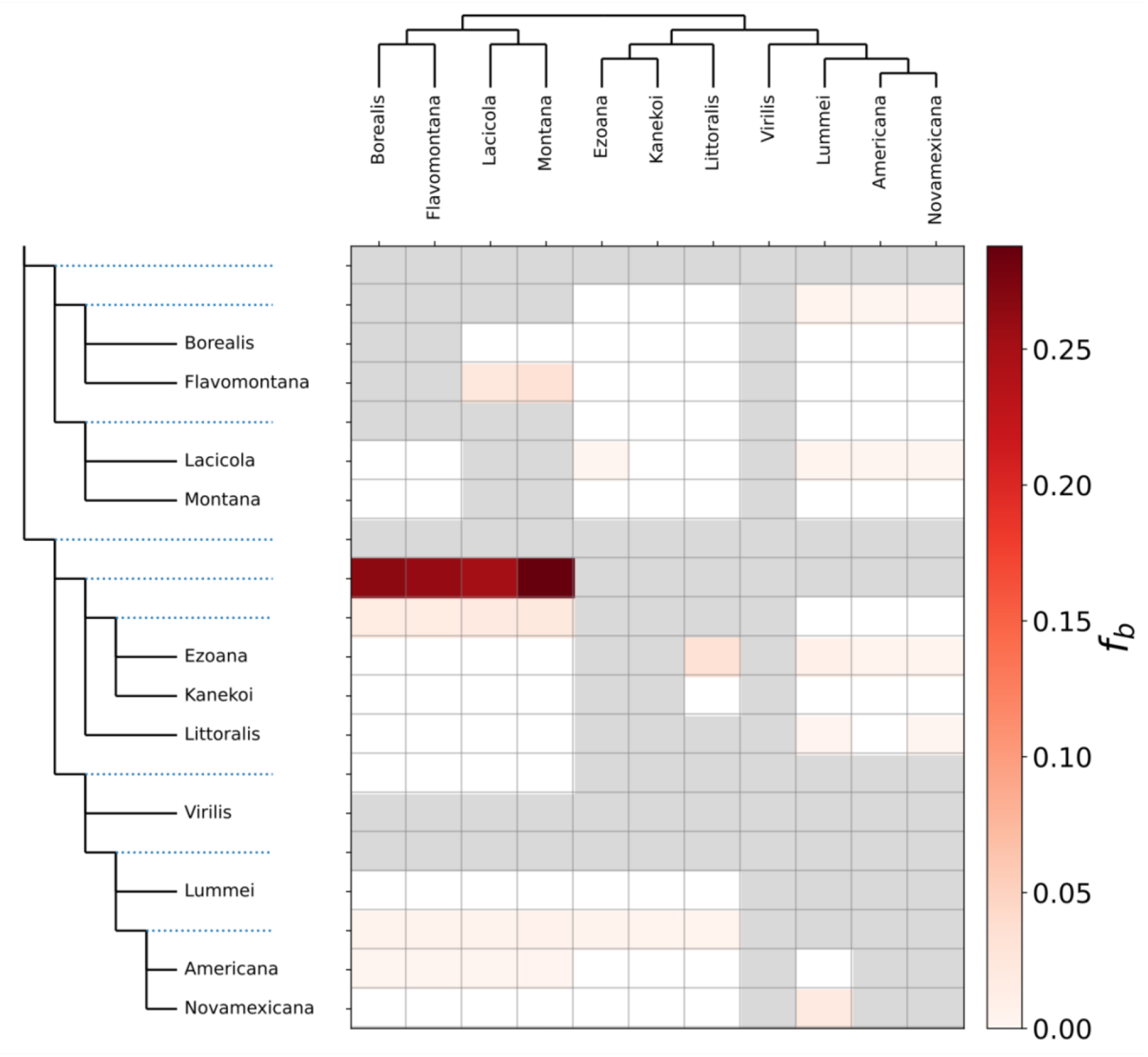
Summary of f-branch (f_b_) tests for introgression in the *virilis* group. Columns represent tips in the phylogeny, whilst rows represent nodes in the tree topology. Colours in cells denote f_b_ statistic between tree nodes and tree tips. Grey cells denote instances where comparisons could not be made.

**Supplementary figure 4:**
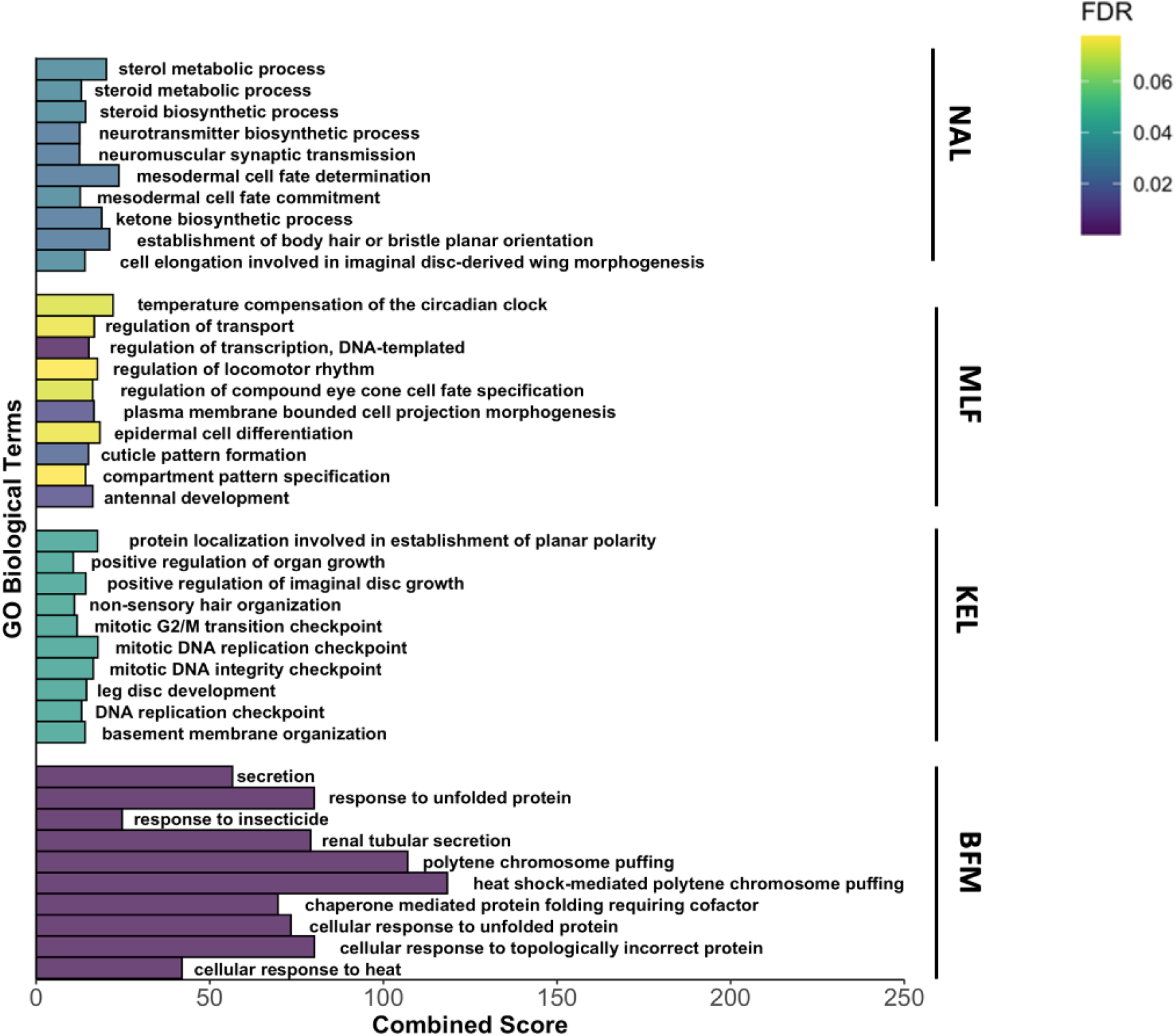
Gene ontology for genes showing signatures of admixture between trios. Here, we show gene ontology biological terms against combined score statistic, where the combined score statistic is calculated by multiplying the log of the p-value for the Fisher exact test by a Z-score computed by assessing deviation from an expected rank, using FlyEnrichr. Biological terms are split by trio, with NAL indicating gene flow from *D. lummei* into either *D. novamexicana* and *D. americana*, MLF indicating gene flow from *D. flavomontana* into *D. montana* and *D. lummei*, KEL indicating gene flow from *D. littoralis* into *D. ezoana* and *D. kanekoi*, and finally BFM indicating gene flow from *D. montana* into *D. flavomontana* and *D. borealis*.

**Supplementary figure 5:**
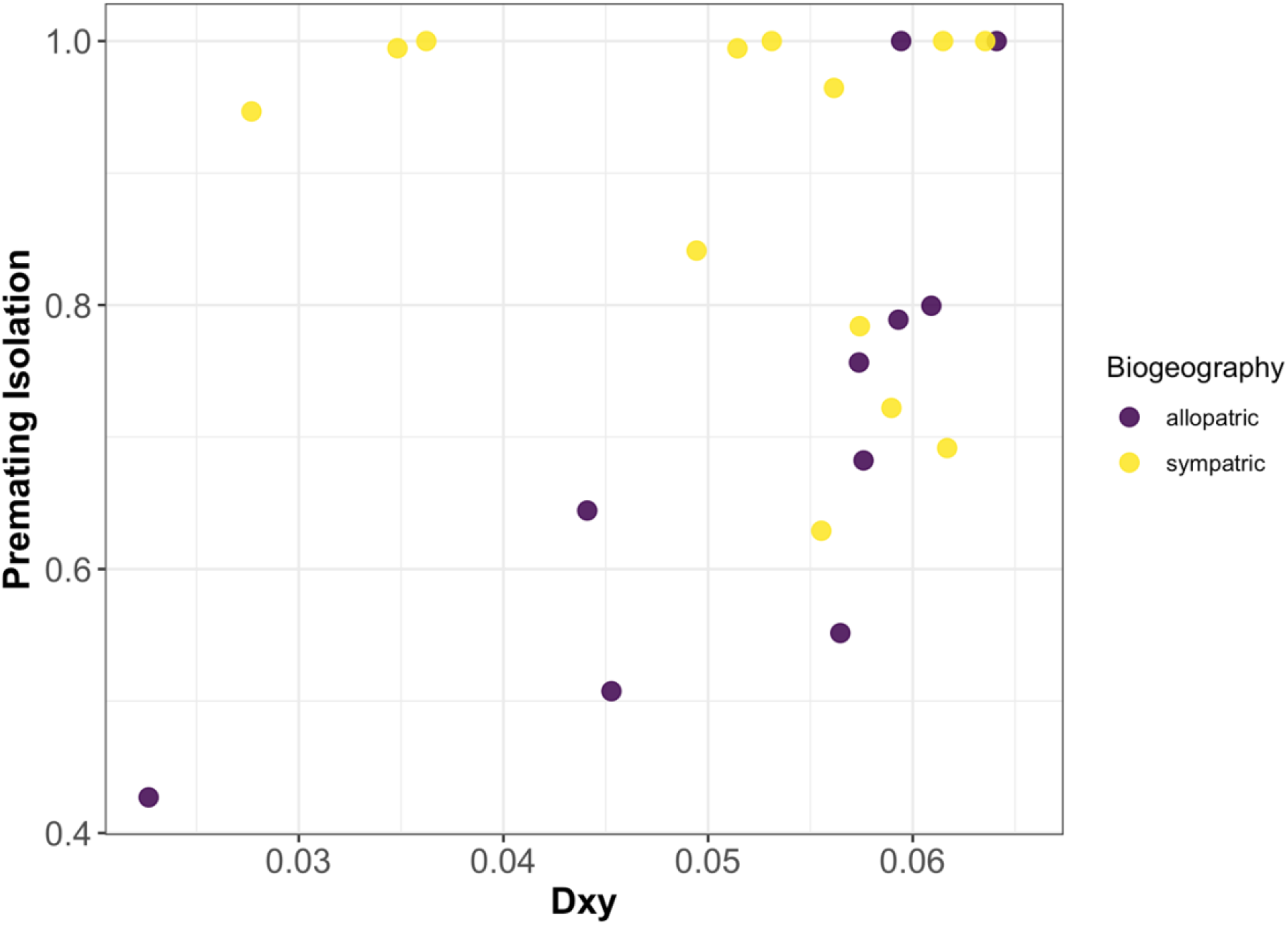
Absolute genetic divergence against pre-mating isolation for species pairs across the *virilis* group. Colours denote species pairs in allopatry (purple) and sympatry (yellow). Estimates of pre-mating isolation were taken from Yukilevich (2014), with 1 denoting complete pre-mating isolation and vice versa.

**Supplementary figure 6:**
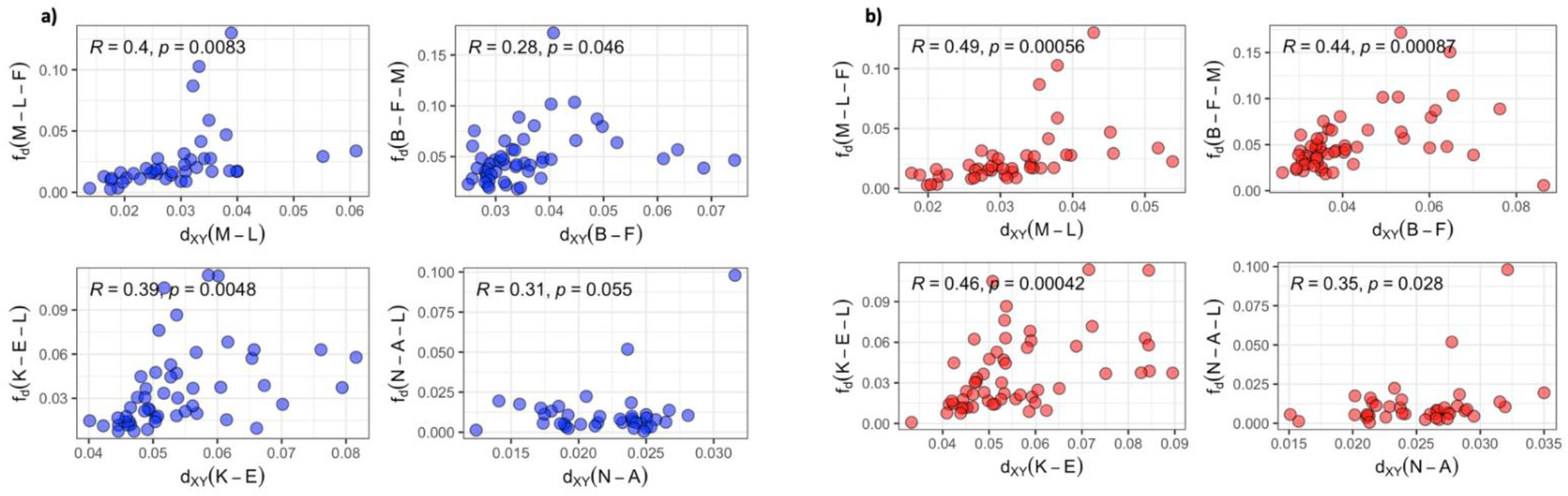
Mean admixture proportions for every scaffold was plotted for closely-related trios across the *virilis* group, against absolute genetic divergence in ‘true’ species pairs. A) Showing the correlation between admixture proportions and genetic divergence in coding regions. B) Showing the correlation between admixture proportions and genetic divergence in non-coding regions. For each plot, Pearson correlation coefficient was calculated.

## References

Abbott, R., D. Albach, S. Ansell, J. W. Arntzen, S. J. E. Baird, N. Bierne, J. Boughman, A. Brelsford, C. A. Buerkle, R. Buggs, R. K. Butlin, U. Dieckmann, F. Eroukhmanoff, A. Grill, S. H. Cahan, J. S. Hermansen, G. Hewitt, A. G. Hudson, C. Jiggins, J. Jones, B. Keller, T. Marczewski, J. Mallet, P. Martinez-Rodriguez, M. Möst, S. Mullen, R. Nichols, A. W. Nolte, C. Parisod, K. Pfennig, A. M. Rice, M. G. Ritchie, B. Seifert, C. M. Smadja, R. Stelkens, J. M. Szymura, R. Väinölä, J. B. W. Wolf, and D. Zinner. 2013. Hybridization and speciation. J. Evol. Biol. 26:229–246.

Ahmed-Braimah, Y. H. 2016. Multiple Genes Cause Postmating Prezygotic Reproductive Isolation in the Drosophila virilis Group. G3. 6:4067–4076.

Ahmed-Braimah, Y. H., and A. L. Sweigart. 2015. A single gene causes an interspecific difference in pigmentation in Drosophila. Genetics 200:331–342.

Andrews, S. 2010. FastQC. Babraham Bioinformatics.

Andrianov, B., I. Goryacheva, N. Mugue, S. Sorokina, T. Gorelova, and V. Mitrofanov. 2010. Comparative analysis of the mitochondrial genomes in Drosophila virilis species group (Diptera: Drosophilidae). Trends Evol. Biol. 2:22–31.

Aspi, A. J., J. Lumme, A. Hoikkala, E. Heikkinen, J. Aspi, J. Lumme, A. Hoikkala, and E. Heikkinen. 1993. Reproductive ecology of the boreal riparian guild of Drosophila. Ecography. 16:65–72.

Bartoli, G., M. Sarnthein, M. Weinelt, H. Erlenkeuser, D. Garbe-Schönberg, and D. W. Lea. 2005. Final closure of Panama and the onset of northern hemisphere glaciation. Earth Planet. Sci. Lett. 237:33–44

Barton, N., and B. O. Bengtsson. 1986. The barrier to genetic exchange between hybridising populations. Heredity (Edinb). 57:357–376.

Brand, C. L., S. B. Kingan, L. Wu, and D. Garrigan. 2013. A selective sweep across species boundaries in Drosophila. Mol. Biol. Evol. 30:2177–2186.

Bubliy, O. A., K. S. Tcheslavskaia, A. M. Kulikov, O. E. Lazebny, and V. G. Mitrofanov. 2007. Variation of wing shape in the Drosophila virilis species group (Diptera: Drosophilidae). J. Zool. Syst. Evol. Res. 46: 38–47

Caletka, B. C., and B. F. McAllister. 2004. A genealogical view of chromosomal evolution and species delimitation in the Drosophila virilis species subgroup. Mol. Phylogenet. Evol. 33:664–670.

Capella-Gutiérrez, S., J. M. Silla-Martínez, and T. Gabaldón. 2009. trimAl: A tool for automated alignment trimming in large-scale phylogenetic analyses. Bioinformatics 25:1972–1973.

Charlesworth, B., J. L. Campos, and B. C. Jackson. 2018. Faster-X evolution: Theory and evidence from Drosophila. Mol. Ecol. 27:3753–3771.

Charlesworth, B., J. A. Coyne, and N. H. Barton. 1987. The Relative Rates of Evolution of Sex Chromosomes and Autosomes. Am. Nat. 130:113–146.

Chekunova, A. I., A. M. Kulikov, S. S. Mikhailovskii, O. E. Lazebny, I. V. Lazebnaya, and V. G. Mitrofanov. 2008. The relationships among the species of the Drosophila virilis group inferred from the gene Ras1 sequences. Russ. J. Genet. 44:286–294.

Chung, H., D. W. Loehlin, H. D. Dufour, K. Vaccarro, J. G. Millar, and S. B. Carroll. 2014. A single gene affects both ecological divergence and mate choice in Drosophila. Science (80-.). 343:1148–1151.

Conesa, A., S. Götz, J. M. García-Gómez, J. Terol, M. Talón, and M. Robles. 2005. Blast2GO: A universal tool for annotation, visualization and analysis in functional genomics research. Bioinformatics. 21(18):3674–6

Coyne, J. A., and H. A. Orr. 1997. “PATTERNS OF SPECIATION IN DROSOPHILA” REVISITED. Evolution (N. Y). 51:295–303.

Coyne, J. A., and H. A. Orr. 1989. Patterns of Speciation in Drosophila. Evolution (N. Y). 43:362–381

Cruickshank, T. E., and M. W. Hahn. 2014. Reanalysis suggests that genomic islands of speciation are due to reduced diversity, not reduced gene flow. Mol. Ecol. 23:3133–3157.

Dasmahapatra, K. K., J. R. Walters, A. D. Briscoe, J. W. Davey, A. Whibley, N. J. Nadeau, A. V. Zimin, C. Salazar, L. C. Ferguson, S. H. Martin, J. J. Lewis, S. Adler, S. J. Ahn, D. A. Baker, S. W. Baxter, N. L. Chamberlain, C. Ritika, B. A. Counterman, T. Dalmay, L. E. Gilbert, K. Gordon, D. G. Heckel, H. M. Hines, K. J. Hoff, P. W. H. Holland, E. Jacquin-Joly, F. M. Jiggins, R. T. Jones, D. D. Kapan, P. Kersey, G. Lamas, D. Lawson, D. Mapleson, L. S. Maroja, A. Martin, S. Moxon, W. J. Palmer, R. Papa, A. Papanicolaou, Y. ick Pauchet, D. A. Ray, N. Rosser, S. L. Salzberg, M. A. Supple, A. Surridge, A. Tenger-Trolander, H. Vogel, P. A. Wilkinson, D. Wilson, J. A. Yorke, F. Yuan, A. L. Balmuth, C. Eland, K. Gharbi, M. Thomson, R. A. Gibbs, Y. Han, J. C. Jayaseelan, C. Kovar, T. Mathew, D. M. Muzny, F. Ongeri, L. L. Pu, J. Qu, R. L. Thornton, K. C. Worley, Y. Q. Wu, M. Linares, M. L. Blaxter, R. H. ffrench-Constant, M. Joron, M. R. Kronforst, S. P. Mullen, R. D. Reed, S. E. Scherer, S. Richards, J. Mallet, W. O. Mc Millan, and C. D. Jiggins. 2012. Butterfly genome reveals promiscuous exchange of mimicry adaptations among species. Nature. 487:94–98

Duranton, M., F. Allal, S. Valière, O. Bouchez, F. Bonhomme, and P. Gagnaire. 2020. The contribution of ancient admixture to reproductive isolation between European sea bass lineages. Evol. Lett. 4:226–242.

Edelman, N. B., P. B. Frandsen, M. Miyagi, B. Clavijo, J. Davey, R. B. Dikow, G. García-accinelli, S. M. Van Belleghem, and N. Patterson. 2019. Genomic architecture and introgression shape a butterfly radiation. Science. (80-.). 599:594–599.

Ellegren, H., L. Smeds, R. Burri, P. I. Olason, N. Backström, T. Kawakami, A. Künstner, H. Mäkinen, K. Nadachowska-Brzyska, A. Qvarnström, S. Uebbing, and J. B. W. Wolf. 2012. The genomic landscape of species divergence in Ficedula flycatchers. Nature. 491:756–760.

Emms, D. M., and S. Kelly. 2019. OrthoFinder: Phylogenetic orthology inference for comparative genomics. Genome Biol. 20:238

Emms, D. M., and S. Kelly. 2015. OrthoFinder: solving fundamental biases in whole genome comparisons dramatically improves orthogroup inference accuracy. Genome Biol. 16:157

Evgen’ev, M. B., H. Zelentsova, H. Poluectova, G. T. Lyozin, V. Veleikodvorskaja, K. I. Pyatkov, L. A. Zhivotovsky, and M. G. Kidwell. 2000. Mobile elements and chromosomal evolution in the virilis group of Drosophila. Proc. Natl. Acad. Sci. 97 (21) 11337-11342

Ferreira, M. S., M. R. Jones, C. M. Callahan, L. Farelo, Z. Tolesa, F. Suchentrunk, P. Boursot, L. S. Mills, P. C. Alves, J. M. Good, and J. Melo-Ferreira. 2020. The Legacy of Recurrent Introgression during the Radiation of Hares. Syst. Biol. 0:1–15.

Flouri, T., X. Jiao, B. Rannala, and Z. Yang. 2018. Species tree inference with BPP using genomic sequences and the multispecies coalescent. Mol. Biol. Evol. 35:2585–2593.

Fonseca, N. A., R. Morales-Hojas, M. Reis, H. Rocha, C. P. Vieira, V. Nolte, C. Schlötterer, and J. Vieira. 2013. Drosophila americana as a model species for comparative studies on the molecular basis of phenotypic variation. Genome Biol. Evol. 5(4):661–79

Fontaine, M. C., J. B. Pease, A. Steele, R. M. Waterhouse, D. E. Neafsey, I. V Sharakhov, X. Jiang, A. B. Hall, F. Catteruccia, E. Kakani, S. N. Mitchell, Y. Wu, H. A. Smith, R. R. Love, M. K. Lawniczak, M. A. Slotman, S. J. Emrich, M. W. Hahn, and N. J. Besansky. 2015. Extensive introgression in a malaria vector species complex revealed by phylogenomics. Science. (80-.). 347:1258524.

Garlovsky, M. D., and R. R. Snook. 2018. Persistent postmating, prezygotic reproductive isolation between populations. Ecol. Evol. 8:9062–9073.

Garlovsky, M. D., L. H. Yusuf, M. G. Ritchie, and R. R. Snook. 2020. Within-population sperm competition intensity does not predict asymmetry in conpopulation sperm precedence. Philos. Trans. R. Soc. Lond. B. Biol. Sci. 375.

Garrigan, D., S. B. Kingan, A. J. Geneva, P. Andolfatto, A. G. Clark, K. R. Thornton, and D. C. Presgraves. 2012. Genome sequencing reveals complex speciation in the Drosophila simulans clade. Genome Res. 22:1499–1511.

Green, R. E., J. Krause, A. W. Briggs, T. Maricic, U. Stenzel, M. Kircher, N. Patterson, H. Li, W. Zhai, M. H.-Y. Fritz, N. F. Hansen, E. Y. Durand, A.-S. Malaspinas, J. D. Jensen, T. Marques-Bonet, C. Alkan, K. Prüfer, M. Meyer, H. A. Burbano, J. M. Good, R. Schultz, A. Aximu-Petri, A. Butthof, B. Höber, B. Höffner, M. Siegemund, A. Weihmann, C. Nusbaum, E. S. Lander, C. Russ, N. Novod, J. Affourtit, M. Egholm, C. Verna, P. Rudan, D. Brajkovic, Z. Kucan, I. Gusic, V. B. Doronichev, L. V Golovanova, C. Lalueza-Fox, M. de la Rasilla, J. Fortea, A. Rosas, R. W. Schmitz, P. L. F. Johnson, E. E. Eichler, D. Falush, E. Birney, J. C. Mullikin, M. Slatkin, R. Nielsen, J. Kelso, M. Lachmann, D. Reich, and S. Pääbo. 2010. A draft sequence of the Neandertal genome. Science. 328:710– 22.

Haddrill, P. R., B. Charlesworth, D. L. Halligan, and P. Andolfatto. 2005. Patterns of intron sequence evolution in Drosophila are dependent upon length and GC content. Genome Biol. 6(8):1–8.

Halligan, D. L., and P. D. Keightley. 2006. Ubiquitous selective constraints in the Drosophila genome revealed by a genome-wide interspecies comparison. Genome Res. 16:875–884.

Han, F., S. Lamichhaney, B. R. Grant, P. R. Grant, L. Andersson, and M. T. Webster. 2017. Gene flow, ancient polymorphism, and ecological adaptation shape the genomic landscape of divergence among Darwin’ s finches. Genome Res. 2017. 27: 1004-1015.

Hoikkala, A., and J. Lumme. 1987. THE GENETIC BASIS OF EVOLUTION OF THE MALE COURTSHIP SOUNDS IN THE DROSOPHILA VIRILIS GROUP. Evolution (N. Y). 41:827–845.

Huerta-Cepas, J., F. Serra, and P. Bork. 2016. ETE 3: Reconstruction, Analysis, and Visualization of Phylogenomic Data. Mol. Biol. Evol. 33:1635–1638.

Jennings, J. H., W. J. Etges, T. Schmitt, and A. Hoikkala. 2014. Cuticular hydrocarbons of Drosophila montana: Geographic variation, sexual dimorphism and potential roles as pheromones. J. Insect Physiol. 61:16–24.

Kalyaanamoorthy, S., B. Q. Minh, T. K. F. Wong, A. Von Haeseler, and L. S. Jermiin. 2017. ModelFinder: Fast model selection for accurate phylogenetic estimates. Nat. Methods 14:587–589.

Katoh, K., and D. M. Standley. 2013. MAFFT multiple sequence alignment software version 7: Improvements in performance and usability. Mol. Biol. Evol. 30(4):772–80

Korunes, K. L., C. A. Machado, and M. A. F. Noor. 2021. Inversions shape the divergence of Drosophila pseudoobscura and Drosophila persimilis on multiple timescales. Evolution (N. Y). 75:1820–1834.

Kuleshov, M. V., J. E. L. Diaz, Z. N. Flamholz, A. B. Keenan, A. Lachmann, M. L. Wojciechowicz, R. L. Cagan, and A. Ma’ayan. 2019. modEnrichr: a suite of gene set enrichment analysis tools for model organisms. Nucleic Acids Res. 47(W1):W183–90.

Kulikov, A. M., A. I. Melnikov, N. G. Gornostaev, O. E. Lazebny, and V. G. Mitrofanov. 2004. Morphological analysis of male mating organ in the Drosophila virilis species group: A multivariate approach. J. Zool. Syst. Evol. Res. 42:135–144.

Kumar, S., A. J. Filipski, F. U. Battistuzzi, S. L. Kosakovsky Pond, and K. Tamura. 2012. Statistics and truth in phylogenomics. Mol. Biol. Evol. 29(2):457–72.

Lamb, A. M., Z. Wang, P. Simmer, H. Chung, and P. J. Wittkopp. 2020. ebony Affects Pigmentation Divergence and Cuticular Hydrocarbons in Drosophila americana and D. novamexicana. Front. Ecol. Evol. 8:1–23.

Lamichhaney, S., J. Berglund, M. S. Almén, K. Maqbool, M. Grabherr, A. Martinez-Barrio, M. Promerová, C.-J. Rubin, C. Wang, N. Zamani, B. R. Grant, P. R. Grant, M. T. Webster, and L. Andersson. 2015. Evolution of Darwin’s finches and their beaks revealed by genome sequencing. Nature. 518:371–375.

Lamichhaney, S., F. Han, M. T. Webster, L. Andersson, B. R. Grant, and P. R. Grant. 2018. Rapid hybrid speciation in Darwin’s finches. Science. (80-.). 359:224–228.

Liimatainen, J. O., and A. Hoikkala. 1998. Interactions of the Males and Females of Three Sympatric Drosophila virilis-Group Species, D. montana, D. littoralis, and D. lummei, (Diptera: Drosophilidae) in Intra- and Interspecific Courtships in the Wild and in the Laboratory. J. Insect Behav. 11(3):399–417

Lohse, K., M. Clarke, M. G. Ritchie, and W. J. Etges. 2015. Genome-wide tests for introgression between cactophilic Drosophila implicate a role of inversions during speciation. Evolution (N. Y). 69:1178–1190.

Lyne, R., R. Smith, K. Rutherford, M. Wakeling, A. Varley, F. Guillier, H. Janssens, W. Ji, P. Mclaren, P. North, D. Rana, T. Riley, J. Sullivan, X. Watkins, M. Woodbridge, K. Lilley, S. Russell, M. Ashburner, K. Mizuguchi, and G. Micklem. 2007. FlyMine: An integrated database for Drosophila and Anopheles genomics. Genome Biol. 8(7):1–6.

Mai, D., M. J. Nalley, and D. Bachtrog. 2020. Patterns of Genomic Differentiation in the Drosophila nasuta Species Complex. Mol. Biol. Evol. 37:208–220.

Malinsky, M., R. J. Challis, A. M. Tyers, S. Schiffels, Y. Terai, B. P. Ngatunga, E. A. Miska, R. Durbin, M. J. Genner, and G. F. Turner. 2015. Genomic islands of speciation separate cichlid ecomorphs in an East African crater lake. Science. (80-.). 350:1493–1498.

Malinsky, M., M. Matschiner, and H. Svardal. 2020. Dsuite -Fast D-statistics and related admixture evidence from VCF files. Mol. Ecol. Resour. 21(2):584–95.

Malinsky, M., H. Svardal, A. M. Tyers, E. A. Miska, M. J. Genner, G. F. Turner, and R. Durbin. 2018. Whole-genome sequences of Malawi cichlids reveal multiple radiations interconnected by gene flow. *Nat*. Ecol. Evol. 2:1940–1955.

Marincovich, L., and A. Y. Gladenkov. 1999. Evidence for an early opening of the Bering Strait. Nature. 397(6715):149–51

Marques, D. A., K. Lucek, J. I. Meier, and S. Mwaiko. 2016. Genomics of Rapid Incipient Speciation in Sympatric Threespine Stickleback. PLoS Genet. 12(2): e1005887.

Marques, D. A., K. Lucek, V. C. Sousa, L. Excoffier, and O. Seehausen. 2019. Admixture between old lineages facilitated contemporary ecological speciation in Lake Constance stickleback. Nat. Commun. 10(1):1–4

Marques, D. A., J. I. Meier, and O. Seehausen. 2019. A Combinatorial View on Speciation and Adaptive Radiation. Trends Ecol. Evol. 34:531–544.

Martin, S. H., and S. M. Van Belleghem. 2017. Exploring Evolutionary Relationships Across the Genome Using Topology Weighting. Genetics. 206:429–438.

Martin, S. H., J. W. Davey, and C. D. Jiggins. 2015. Evaluating the use of ABBA-BABA statistics to locate introgressed loci. Mol. Biol. Evol. 32:244–257.

McGee, M. D., S. R. Borstein, J. I. Meier, D. A. Marques, S. Mwaiko, A. Taabu, M. A. Kishe, B. O’Meara, R. Bruggmann, L. Excoffier, and O. Seehausen. 2020. The ecological and genomic basis of explosive adaptive radiation. Nature. 586(7827):75–9

Meier, J. I., D. A. Marques, S. Mwaiko, C. E. Wagner, L. Excoffier, and O. Seehausen. 2017. Ancient hybridization fuels rapid cichlid fish adaptive radiations. Nat. Commun. 8:1–11.

Meisel, R. P., and T. Connallon. 2013. The faster-X effect: Integrating theory and data. Trends Genet. 29:537–544.

Mendes, F. K., and M. W. Hahn. 2018. Why Concatenation Fails Near the Anomaly Zone. Syst. Biol. 67:158–169.

Mirol, P. M., M. A. Schäfer, L. Orsini, J. Routtu, C. Schlötterer, A. Hoikkala, and R. K. Butlin. 2007. Phylogeographic patterns in Drosophila montana. Mol. Ecol. 16(5):1085–97.

Morales-Hojas, R., M. Reis, C. P. Vieira, and J. Vieira. 2011. Resolving the phylogenetic relationships and evolutionary history of the Drosophila virilis group using multilocus data. Mol. Phylogenet. Evol. 60:249–258.

Nachman, M. W., and B. A. Payseur. 2012. Recombination rate variation and speciation: Theoretical predictions and empirical results from rabbits and mice. Philos. Trans. R. Soc. B Biol. Sci. 367:409–421.

Nadeau, N. J., S. H. Martin, K. M. Kozak, C. Salazar, K. K. Dasmahapatra, J. W. Davey, S. W. Baxter, M. L. Blaxter, J. Mallet, and C. D. Jiggins. 2013. Genome-wide patterns of divergence and gene flow across a butterfly radiation. Mol. Ecol. 22:814–826.

Nelson, T. C., and W. A. Cresko. 2018. Ancient genomic variation underlies repeated ecological adaptation in young stickleback populations. Evol. Lett. 2:9–21.

Noor, M. A. F., and S. M. Bennett. 2009. Islands of speciation or mirages in the desert? Examining the role of restricted recombination in maintaining species. Heredity (Edinb). 103:439–444.

Nosil, P., D. J. Funk, and D. Ortiz-Barrientos. 2009. Divergent selection and heterogeneous genomic divergence. Mol. Ecol. 18(3), 375–402.

Nurminsky, D. I., E. N. Moriyama, E. R. Lozovskaya, and D. L. Hartl. 1996. Molecular phylogeny and genome evolution in the Drosophila virilis species group: Duplications of the alcohol Dehydrogenase gene. Mol. Biol. Evol. 13(1):132–49.

Orr, H. A., and J. A. Coyne. 1989. The genetics of postzygotic isolation in the Drosophila virilis group. Genetics 121:527–537.

Orsini, L., S. Huttunen, and C. Schlötterer. 2004. A multilocus microsatellite phylogeny of the Drosophila virilis group. Heredity (Edinb). 93(2):161–5.

Oziolor, E. M., N. M. Reid, S. Yair, K. M. Lee, S. Guberman VerPloeg, P. C. Bruns, J. R. Shaw, A. Whitehead, and C. W. Matson. 2019. Adaptive introgression enables evolutionary rescue from extreme environmental pollution. Science (80-.). 364:455–457.

Parker, D. J., M. G. Ritchie, and M. Kankare. 2016. Preparing for winter: The transcriptomic response associated with different day lengths in Drosophila montana. G3 6:1373–1381.

Parker, D. J., L. Vesala, M. G. Ritchie, A. Laiho, A. Hoikkala, and M. Kankare. 2015. How consistent are the transcriptome changes associated with cold acclimation in two species of the Drosophila virilis group? Heredity (Edinb). 115:13–21.

Patterson, J. 1952. Revision of the montana complex of the virilis species group. University of Texas Publ. 5204:20–34

Pease, J. B., D. C. Haak, M. W. Hahn, and L. C. Moyle. 2016. Phylogenomics Reveals Three Sources of Adaptive Variation during a Rapid Radiation. PLoS Biol. 14:1–24.

Perri, A. R., K. J. Mitchell, A. Mouton, S. Álvarez-Carretero, A. Hulme-Beaman, J. Haile, A. Jamieson, J. Meachen, A. T. Lin, B. W. Schubert, C. Ameen, E. E. Antipina, P. Bover, S. Brace, A. Carmagnini, C. Carøe, J. A. Samaniego Castruita, J. C. Chatters, K. Dobney, M. dos Reis, A. Evin, P. Gaubert, S. Gopalakrishnan, G. Gower, H. Heiniger, K. M. Helgen, J. Kapp, P. A. Kosintsev, A. Linderholm, A. T. Ozga, S. Presslee, A. T. Salis, N. F. Saremi, C. Shew, K. Skerry, D. E. Taranenko, M. Thompson, M. V. Sablin, Y. V. Kuzmin, M. J. Collins, M. H. S. Sinding, M. T. P. Gilbert, A. C. Stone, B. Shapiro, B. Van Valkenburgh, R. K. Wayne, G. Larson, A. Cooper, and L. A. F. Frantz. 2021. Dire wolves were the last of an ancient New World canid lineage. Nature. 591(7848):87–91

Poikela, N., J. Kinnunen, M. Wurdack, H. Kauranen, T. Schmitt, M. Kankare, R. R. Snook, and A. Hoikkala. 2019. Strength of sexual and postmating prezygotic barriers varies between sympatric populations with different histories and species abundances. Evolution (N. Y). 73:1182–1199.

Purcell, S., B. Neale, K. Todd-Brown, L. Thomas, M. A. R. Ferreira, D. Bender, J. Maller, P. Sklar, P. I. W. De Bakker, M. J. Daly, and P. C. Sham. 2007. PLINK: A tool set for whole-genome association and population-based linkage analyses. Am. J. Hum. Genet. 81(3):559–75

Racimo, F., D. Marnetto, and E. Huerta-Sánchez. 2016. Signatures of archaic adaptive introgression in present-day human populations. Mol. Biol. Evol. 34(2):296–317.

Ravinet, M., R. Faria, R. K. Butlin, J. Galindo, N. Bierne, M. Rafajlović, M. A. F. Noor, M. B, and A. M. Westram. 2017. Interpreting the genomic landscape of speciation : a road map for finding barriers to gene flow. Journal of evolutionary biology. 30:1450–1477.

Reis, M., C. P. Vieira, R. Lata, N. Posnien, and J. Vieira. 2018. Origin and consequences of chromosomal inversions in the virilis group of Drosophila. Genome Biol. Evol. 10:3152– 3166.

Reis, M., G. Wiegleb, J. Claude, R. Lata, B. Horchler, N. T. Ha, C. Reimer, C. P. Vieira, J. Vieira, and N. Posnien. 2020. Multiple loci linked to inversions are associated with eye size variation in species of the Drosophila virilis phylad. Sci. Rep. 10(1):1–7.

Ritchie, M. G., M. Saarikettu, S. Livingstone, and A. Hoikkala. 2001. Characterisation of female preference functions for Drosophila montana courtship song and a test of the temperature coupling hypothesis. Evolution (N. Y). 55:721–727.

Robinson, M. M., H. J. Dowsett, and M. A. Chandler. 2008. Pliocene role in assessing future climate impacts. Eos (Washington. DC). 89(49):501–2

Roux, C., C. Fraïsse, J. Romiguier, Y. Anciaux, N. Galtier, and N. Bierne. 2016. Shedding Light on the Grey Zone of Speciation along a Continuum of Genomic Divergence. PLoS Biol. 14:1–22.

Ruggieri, E., T. Herbert, K. T. Lawrence, and C. E. Lawrence. 2009. Change point method for detecting regime shifts in paleoclimatic time series: Application to δ18O time series of the Plio-Pleistocene. Paleoceanography. 24(1).

Salichos, L., and A. Rokas. 2013. Inferring ancient divergences requires genes with strong phylogenetic signals. Nature 497:327–331.

Salminen, T. S., L. Vesala, A. Laiho, M. Merisalo, A. Hoikkala, and M. Kankare. 2015. Seasonal gene expression kinetics between diapause phases in Drosophila virilis group species and overwintering differences between diapausing and non-diapausing females. Sci. Rep. 5:1–13.

Samuk, K., G. L. Owens, K. E. Delmore, S. E. Miller, D. J. Rennison, and D. Schluter. 2017. Gene flow and selection interact to promote adaptive divergence in regions of low recombination. Mol. Ecol. 26:4378–4390.

Sankararaman, S., S. Mallick, N. Patterson, and D. Reich. 2016. The Combined Landscape of Denisovan and Neanderthal Ancestry in Present-Day Humans. Curr. Biol. 26:1241–1247.

Sankararaman, S., N. Patterson, D. Reich, S. Mallick, M. Dannemann, K. Prüfer, J. Kelso, S. Pääbo, N. Patterson, and D. Reich. 2014. The genomic landscape of Neanderthal ancestry in present-day humans. Nature. 507:354–7.

Schrider DR, Ayroles J, Matute DR, Kern AD (2018) Supervised machine learning reveals introgressed loci in the genomes of Drosophila simulans and D. sechellia. PLoS Genet 14(4): e1007341.

Small, S. T., F. Labbé, N. F. Lobo, L. L. Koekemoer, C. H. Sikaala, D. E. Neafsey, M. W. Hahn, M. C. Fontaine, and N. J. Besansky. 2020. Radiation with reticulation marks the origin of a major malaria vector. Proc. Natl. Acad. Sci. 117(50):31583–90

Stankowski S, Chase MA, Fuiten AM, Rodrigues MF, Ralph PL, Streisfeld MA (2019) Widespread selection and gene flow shape the genomic landscape during a radiation of monkeyflowers. PLoS Biol 17(7): e3000391.

Anton Suvorov, Bernard Y. Kim, Jeremy Wang, Ellie E. Armstrong, David Peede, Emmanuel R.R. D’Agostino, Donald K. Price, Peter Waddell, Michael Lang, Virginie Courtier-Orgogozo, Jean R. David, Dmitri Petrov, Daniel R. Matute, Daniel R. Schrider, Aaron A. Comeault (2021). Widespread introgression across a phylogeny of 155 Drosophila genomes. Current Biology.

Svardal, H., F. X. Quah, M. Malinsky, B. P. Ngatunga, E. A. Miska, W. Salzburger, M. J. Genner, G. F. Turner, and R. Durbin. 2020. Ancestral hybridization facilitated species diversification in the lake malawi cichlid fish adaptive radiation. Mol. Biol. Evol. 37:1100–1113.

Sweigart, A. L. 2010. The genetics of postmating, prezygotic reproductive isolation between Drosophila virilis and D. americana. Genetics 184:401–410.

Thawornwattana, Y., D. Dalquen, and Z. Yang. 2018. Coalescent analysis of phylogenomic data confidently resolves the species relationships in the Anopheles gambiae species complex. Mol. Biol. Evol. 35:2512–2527.

Throckmorton, H. L. 1982. The virilis Species group. Genet Bioogy Drosoph. 3:227–96.

Turissini, D. A., and D. R. Matute. 2017. Fine scale mapping of genomic introgressions within the Drosophila yakuba clade. PLoS genetics. 13(9):e1006971

Tyukmaeva, V. I., P. Veltsos, J. Slate, E. Gregson, H. Kauranen, M. Kankare, M. G. Ritchie, R. K. Butlin, and A. Hoikkala. 2015. Localization of quantitative trait loci for diapause and other photoperiodically regulated life history traits important in adaptation to seasonally varying environments. Mol. Ecol. 24:2809–2819.

Valencia-Montoya, W. A., S. Elfekih, H. L. North, J. I. Meier, I. A. Warren, W. T. Tay, K. H. J. Gordon, A. Specht, S. V. Paula-Moraes, R. Rane, T. K. Walsh, and C. D. Jiggins. 2020. Adaptive introgression across semipermeable species boundaries between local helicoverpa zea and invasive helicoverpa armigera moths. Mol. Biol. Evol. 37:2568–2583.

Veltsos, P., C. Wicker-Thomas, R. K. Butlin, A. Hoikkala, and M. G. Ritchie. 2012. Sexual selection on song and cuticular hydrocarbons in two distinct populations of Drosophila montana. Ecol. Evol. 2:80–94.

Vernot, B., and J. M. Akey. 2014. Resurrecting Surviving Neandeltal Linages from Modern Human Genomes. Science (80-.). 343:1017–1021.

Vesala, L., T. S. Salminen, A. Laiho, A. Hoikkala, and M. Kankare. 2012. Cold tolerance and cold-induced modulation of gene expression in two Drosophila virilis group species with different distributions. Insect Mol. Biol. 21:107–118.

Wang, B. cheng, J. Park, H. aki Watabe, J. jun Gao, J. gong Xiangyu, T. Aotsuka, H. wei Chen, and Y. ping Zhang. 2006. Molecular phylogeny of the Drosophila virilis section (Diptera: Drosophilidae) based on mitochondrial and nuclear sequences. Mol. Phylogenet. Evol. 40(2):484–500.

Whitney, K. D., R. A. Randell, and L. H. Rieseberg. 2010. Adaptive introgression of abiotic tolerance traits in the sunflower Helianthus annuus. New Phytol. 187(1):230–9.

Whitney, K. D., R. A. Randell, and L. H. Rieseberg. 2006. Adaptive introgression of herbivore resistance traits in the weedv sunflower Helianthus annuus. Am. Nat. 167(6):794–807.

Wiberg, R. A. W., V. Tyukmaeva, A. Hoikkala, M. G. Ritchie, and M. Kankare. 2021. Cold adaptation drives population genomic divergence in the ecological specialist, Drosophila montana. Mol. Ecol.

Wittkopp, P. J., J. R. True, and S. B. Carroll. 2002. Reciprocal functions of the Drosophila Yellow and Ebony proteins in the development and evolution of pigment patterns. Development 129:1849–1858.

Wittkopp, P. J., K. Vaccaro, and S. B. Carroll. 2002. Evolution of yellow gene regulation and pigmentation in Drosophila. Curr. Biol. 12:1547–1556.

Wolf, J. B. W., and H. Ellegren. 2017. Making sense of genomic islands of differentiation in light of speciation. Nat. Rev. Genet. 18:87–100.

Wu, C. I., & Ting, C. T. (2004). Genes and speciation. Nat. Rev. Genet. 5(2), 114–122.

Yang, Z., and R. Nielsen. 2002. Codon-Substitution Models for Detecting Molecular Adaptation at Individual Sites Along Specific Lineages. Mol. Biol. Evol. 19:908–917.

Yang, Z., R. Nielsen, N. Goldman, and A. M. K. Pedersen. 2000. Codon-substitution models for heterogeneous selection pressure at amino acid sites. Genetics. 155(1):431–49.

Yang, Z., and T. Zhu. 2018. Bayesian selection of misspecified models is overconfident and may cause spurious posterior probabilities for phylogenetic trees. Proc. Natl. Acad. Sci. U. S. A. 115(8):1854–9.

Yukilevich, R. 2014. The rate test of speciation: Estimating the likelihood of non-allopatric speciation from reproductive isolation rates in drosophila. Evolution (N. Y). 68:1150– 1162.

Zhang, W., K. K. Dasmahapatra, J. Mallet, G. R. P. Moreira, and M. R. Kronforst. 2016. Genome-wide introgression among distantly related Heliconius butterfly species. Genome Biol. 17(1):1–5.

Zhao, L., and W. a Jones. 2012. Expression of heat shock protein genes in insect stress responses. Invertebr. Surviv. J. 9:93–101.

Zimin, A. V., G. Marçais, D. Puiu, M. Roberts, S. L. Salzberg, and J. A. Yorke. 2013. The MaSuRCA genome assembler. Bioinformatics 29:2669–2677.

